# Increasing prevalence of plant-fungal symbiosis across two centuries of environmental change

**DOI:** 10.1101/2024.09.05.611506

**Authors:** Joshua C. Fowler, Jacob Moutouama, Tom E. X. Miller

## Abstract

Species’ distributions and abundances are shifting in response to ongoing global climate change. Mutualistic microbial symbionts can provide hosts with protection from environmental stress that may promote resilience under environmental change, however this change may also disrupt species interactions and lead to declines in hosts and/or symbionts. Symbionts preserved within natural history specimens offer a unique opportunity to quantify changes in microbial symbiosis across broad temporal and spatial scales. We asked how the prevalence of seed-transmitted fungal symbionts of grasses (*Epichloë* endophytes) has changed over time in response to climate change, and how these changes vary across host species’ distributions. Specifically, we examined 2,346 herbarium specimens of three grass host species (*Agrostis hyemalis*, *Agrostis perennans*, *Elymus virginicus* ) collected over the past two centuries (1824 – 2019) for the presence or absence of *Epichloë* symbiosis. Analysis of an approximate Bayesian spatially-varying coefficients model revealed that endophytes increased in prevalence over the last two centuries from ca. 25% to ca. 75% prevalence, on average, across three host species. Changes in seasonal climate drivers were associated with increasing endophyte prevalence. Notably, increasing precipitation during the peak growing season for *Agrostis* species and decreasing precipitation for *E. virginicus* were associated with increasing endophyte prevalence. Changes in the variability of precipitation and temperature during off-peak seasons were also important predictors of increasing endophyte prevalence. Our model performed favorably in an out-of-sample predictive test with contemporary survey data from across 63 populations, a rare extra step in collections-based research. However, there was greater local-scale variability in endophyte prevalence in contemporary data compared to model predictions, suggesting new directions that could improve predictive accuracy. Our results provide novel evidence for a cryptic biological response to climate change that may contribute to the resilience of host-microbe symbiosis through fitness benefits to symbiotic hosts.

## Introduction

Understanding how biotic interactions are altered by global change is a major goal of basic and applied ecological research (Blois et al., 2013; Gilman et al., 2010). Documented responses to environmental change, such as shifts in species’ distributions (Aitken et al., 2008) and phenology (Piao et al., 2019), are typically blind to concurrent changes in associated biotic interactions. Empirically evaluating these biotic changes – whether interacting species shift in tandem with their partners or not (HilleRisLambers et al., 2013) – is crucial to predicting the reorganization of Earth’s biodiversity under global change. Such evaluations have been limited because few datasets on species interactions extend over sufficiently long time scales of contemporary climate change (Poisot et al., 2021).

Natural history specimens, which were originally collected to document and preserve taxonomic diversity, present a unique opportunity to explore long-term changes in biodiversity and ecological interactions across broad spatial and temporal scales (Davis, 2023; Meineke et al., 2018). Natural history collections, built and maintained by the efforts of thousands of scientists, are invaluable time machines, primarily comprised of physical specimens of organisms along with information about the time and place of their collection. These specimens often preserve physical legacies of ecological processes and species’ interactions from dynamically changing environments across time and space (Lendemer et al., 2020). For example, previous researchers have examined the flowers, pollen grains, and leaves of specimens within plant collections (herbaria) to document shifts in reproductive phenology (Berg et al., 2019; Park et al., 2019; Willis et al., 2017), pollination (Duan et al., 2019; Pauw and Hawkins, 2011), and herbivory (Meineke et al., 2019) related to anthropogenic climate change. Herbarium specimens have also been used to identify the origins and population genomics of plant diseases such as *Phytophora*, the Irish potato famine pathogen (Ristaino et al., 2001; Ristaino, 2002; Yoshida et al., 2013), and have been proposed as vehicles to track other emerging plant pathogens (Bradshaw et al., 2021; Ristaino, 2020). However, few previous studies have leveraged biological collections to examine climate change-related shifts in a particularly common type of interaction: mutualistic microbial symbiosis.

Microbial symbionts are common to all macroscopic organisms and can have important effects on their hosts’ survival, growth and reproduction (McFall-Ngai et al., 2013; Rodriguez et al., 2009). Many microbial symbionts act as mutualists, engaging in reciprocally beneficial interactions with their hosts in ways that can ameliorate environmental stress. For example, bacterial symbionts of insects, such as *Wolbachia*, can improve their hosts’ thermal tolerance (Renoz et al., 2019; Truitt et al., 2019), and arbuscular mycorrhizal fungi, documented in 70-90% of families of land plants (Parniske, 2008), allow their hosts to persist through drought conditions by improving water and nutrient uptake (Cheng et al., 2021). On the other hand, changes in the mean and variance of environmental conditions may disrupt microbial mutualisms by changing the costs and benefits of the interaction for each partner in ways that can cause the interaction to deteriorate (Aslan et al., 2013; Fowler et al., 2024). Coral bleaching (the loss of symbiotic algae) due to temperature stress (Sully et al., 2019) is perhaps the best known example, but this phenomenon is not unique to corals. Lichens exposed to elevated temperatures experienced loss of photosynthetic function along with changes in the composition of their algal symbiont community (Meyer et al., 2022). How commonly and under what conditions microbial mutualisms deteriorate or strengthen under climate change remain unanswered questions (Frederickson, 2017). Previous work suggests that these alternative responses may depend on the intimacy and specialization of the interaction as well as the physiological tolerances of the mutualist partners (Rafferty et al., 2015; Toby Kiers et al., 2010; Warren and Bradford, 2014).

Understanding how microbial symbioses are affected by climate change is additionally complicated by spatial heterogeneity in the direction and magnitude of environmental change (IPCC, 2021). Beneficial symbionts are likely able to shield their hosts from environmental stress in locations that experience a small degree of change, but symbionts in locations that experience changes of large magnitude may be pushed beyond their physiological limits (Webster et al., 2008). Additionally, symbionts are often unevenly distributed across their host’s distribution. Facultative symbionts may be absent from portions of the host range (Afkhami et al., 2014), and hosts may engage with a diversity of partners (different symbiont species or locally-adapted strains) across environments (Fowler et al., 2023; Frade et al., 2008; Rolshausen et al., 2018). Identifying broader spatial trends in symbiont prevalence is therefore an important step in developing predictions for where to expect changes in the symbiosis in future climates.

*Epichloë* fungal endophytes are specialized symbionts of cool-season grasses, documented in ∼ 30% of cool-season grass species (Leuchtmann, 1992). They are predominantly transmitted vertically from maternal plants to offspring through seeds. Vertical transmission creates a feedback between the fitness of host and symbiont (Douglas, 1998; Fine, 1975; Rudgers et al., 2009). Over time, endophytes that act as mutualists should rise in prevalence within a host population, particularly under environmental conditions that elicit protective benefits (Donald et al., 2021). *Epichloë* are known to improve their hosts’ drought tolerance (Decunta et al., 2021) and protect their hosts against herbivores (Crawford et al., 2010) and pathogens (Xia et al., 2018) likely through the production of a diverse suite of alkaloids and other secondary metabolites. The fitness feedback induced by vertical transmission leads to the prediction that endophyte prevalence should be high in populations where these fitness benefits are most important. Previous survey studies of contemporary populations have documented large-scale spatial patterns in endophyte prevalence structured by environmental gradients (Afkhami, 2012; Bazely et al., 2007; Granath et al., 2007; Sneck et al., 2017). We predicted that endophyte prevalence should track temporal changes in environmental drivers (i.e. drought) that elicit strong fitness benefits.

Early research on *Epichloë* used herbarium specimens to describe the broad taxonomic diversity of grass host species that harbor these symbionts (White and Cole, 1985), establishing that endophyte symbiosis could be identified in plant tissue from as early as 1851. However, no subsequent studies, to our knowledge, have used the vast resources of biological collections to quantitatively assess spatio-temporal trends in endophyte prevalence and their environmental correlates. Grasses are commonly collected and identified based on the presence of their reproductive structures, meaning that preserved specimens typically contain seeds, conveniently preserving the seed-transmitted fungi along with their host plants on herbarium sheets. This creates the opportunity to leverage the unique spatio-temporal sampling of herbarium collections to examine the response of this symbiosis to historical climate change. However, the predictive ability derived from historical analyses is rarely tested against contemporary data (Lee et al., 2024). Critically evaluating whether insights from historical reconstruction are predictive of variation across contemporary populations is a crucial step for the field to move from reading signatures of the past to forecasting ecological dynamics into the future.

In this study, we assessed the long-term responses of *Epichloë* endophyte symbiosis to climate change through the use of herbarium specimens of three North American host grass species (*Agrostis hyemalis*, *Agrostis perennans*, and *Elymus virginicus* ). We first addressed questions describing spatial and temporal trends in endophyte prevalence: (i) How has endophyte prevalence changed over the past two centuries? and (ii) How spatially variable are temporal trends in endophyte prevalence across eastern North America? We then addressed how climate change may be driving trends in endophyte prevalence by asking: (iii) What is the relationship between temporal trends in endophyte prevalence and associated changes in climate drivers? We predicted that overall endophyte prevalence would increase over time in tandem with climate change, and that localized hotspots of endophyte change would correspond spatially to hotspots of climate warming and drying. Finally, we evaluated (iv) how our model, built on data from historic specimens, performed in an out-of-sample test using data on endophyte prevalence from contemporary surveys of host populations. To answer these questions we examined a total of 2, 346 historic specimens collected across eastern North America between 1824 and 2019, and evaluated model performance against contemporary surveys comprising 1,442 individuals from 63 populations surveyed between 2013 and 2020.

## Methods

### Focal species

Our surveys focused on three native North American grasses: *Agrostis hyemalis*, *Agrostis perennans*, and *Elymus virginicus* that host *Epichloë* symbionts. These cool-season grass species have broad distributions covering much the eastern United States (Fig. 1) and are commonly represented in natural history collections. Cool-season grasses grow during the cooler temperatures of spring and autumn due to their reliance on *C*_3_ photosynthesis. *A. hyemalis* is a small short-lived perennial species that germinates in autumn to late winter winter and typically flowers between March and July (most common collection month: May). *A. perennans* is of similar stature but is longer lived than *Agrostis hyemalis* and flowers in late summer and early autumn (most common collection month: September). *A. perennans* is more sparsely distributed, tending to be found in shadier and moister habitats, while *A. hyemalis* is commonly found in open and recently disturbed habitats. Both *Agrostis* species are recorded from throughout the Eastern US, but *A. perennans* has a slighty more northern distribution, whereas *A. hyemalis* is found rarely as far north as Canada and is listed as a rare plant in Minnesota. *E. virginicus* is a larger and longer-lived species that is more broadly distributed than the *Agrostis* species. It begins flowering as early as March or April but continues throughout the summer (most common collection month: July).

**Figure 1:**
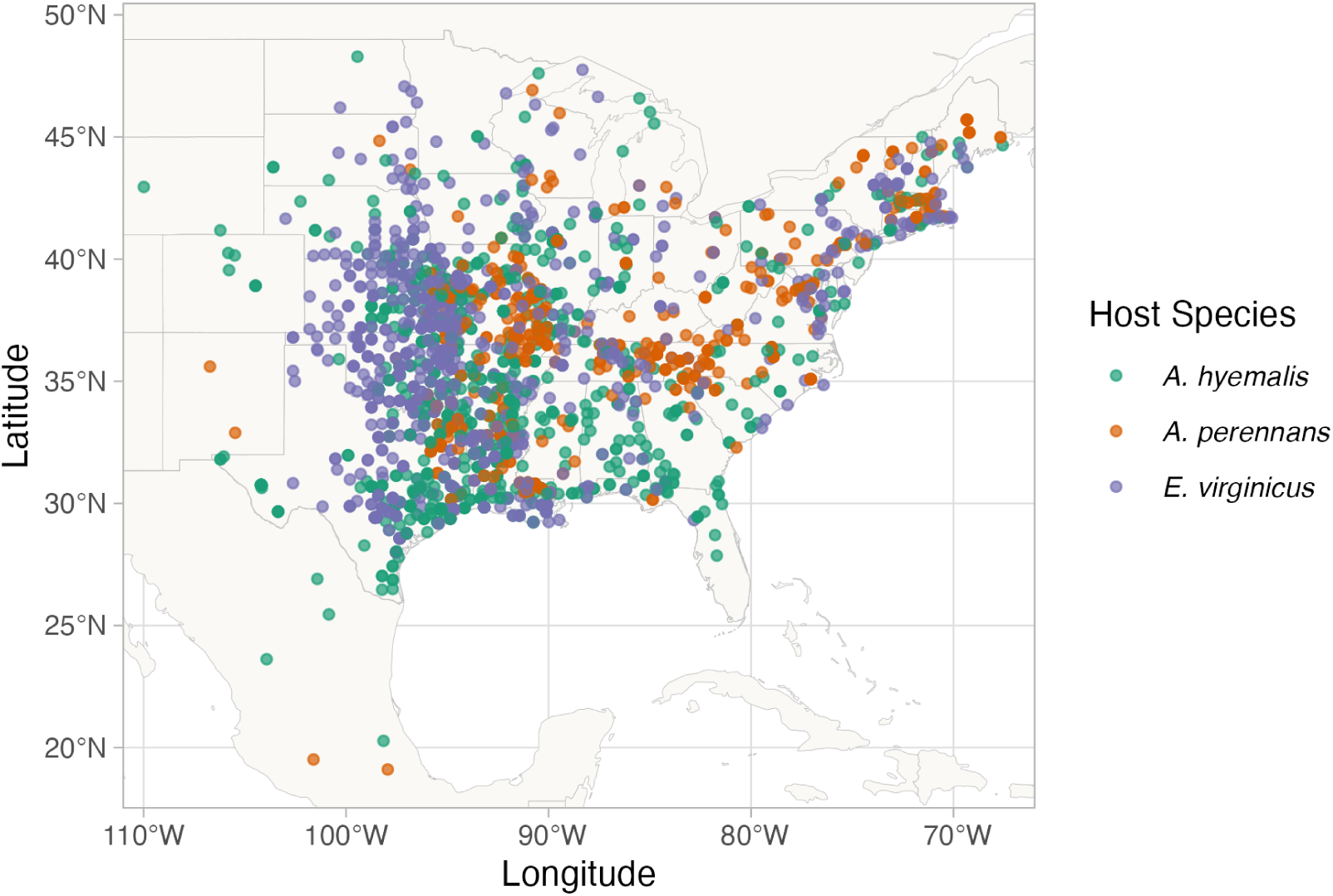
Collection locations of herbarium specimens sampled for *Epichloë* endophytes. Specimens span eastern North America from nine herbaria, and are colored by host species (*A. hyemalis*: green, *A. perennans*: orange, *E. virginicus*: purple). Map lines delineate study areas and do not necessarily depict accepted national boundaries.

Both *Agrostis* species host *Epichloë amarillans* (Craven et al., 2001; Leuchtmann et al., 2014), and *Elymus virginicus* typically hosts *Epichloë elymi* (Clay and Schardl, 2002). The fungal symbionts primarily reproduce asexually and are passed from maternal plant to offspring by vertical transmission through seeds. Some host species have been shown to partner with multiple symbiont species or strains, and in some cases multiple symbiont lineages can co-exist within a host population (Mc Cargo et al., 2014). However, surveys have typically found limited *Epichloë* genotypic diversity within host populations (Treindl et al., 2023). Across host populations, concentrations of biologically-active alkaloids and the genes associated with their production vary substantially (Schardl et al., 2012). In this analysis, we focus on the presence/absence of *Epichloë* symbionts, and we discuss potential implications of symbiont genotypic diversity in the Discussion.

### Herbarium surveys

We visited nine herbaria between 2019 and 2022 (see Table A1 for a summary of specimens included from each collection). With permission from herbarium staff, we acquired seed samples from 1135 *A. hyemalis* specimens collected between 1824 and 2019, 357 *A. perennans* specimens collected between 1863 and 2017, and 854 *E. virginicus* specimens collected between 1839 and 2019 (Fig. 1, Fig. 2A, Fig. A1). We chose our focal species in part because they are commonly represented in herbarium collections and produce many seeds, meaning that small samples would not diminish the value of specimens for future studies. We collected 5-10 seeds per specimen after examining the herbarium sheet under a dissecting microscope to ensure that we collected mature seeds, not florets or unfilled seeds, fit for our purpose of identifying fungal endophytes with microscopy. We excluded specimens for which information about the collection location and date were unavailable.

**Figure 2:**
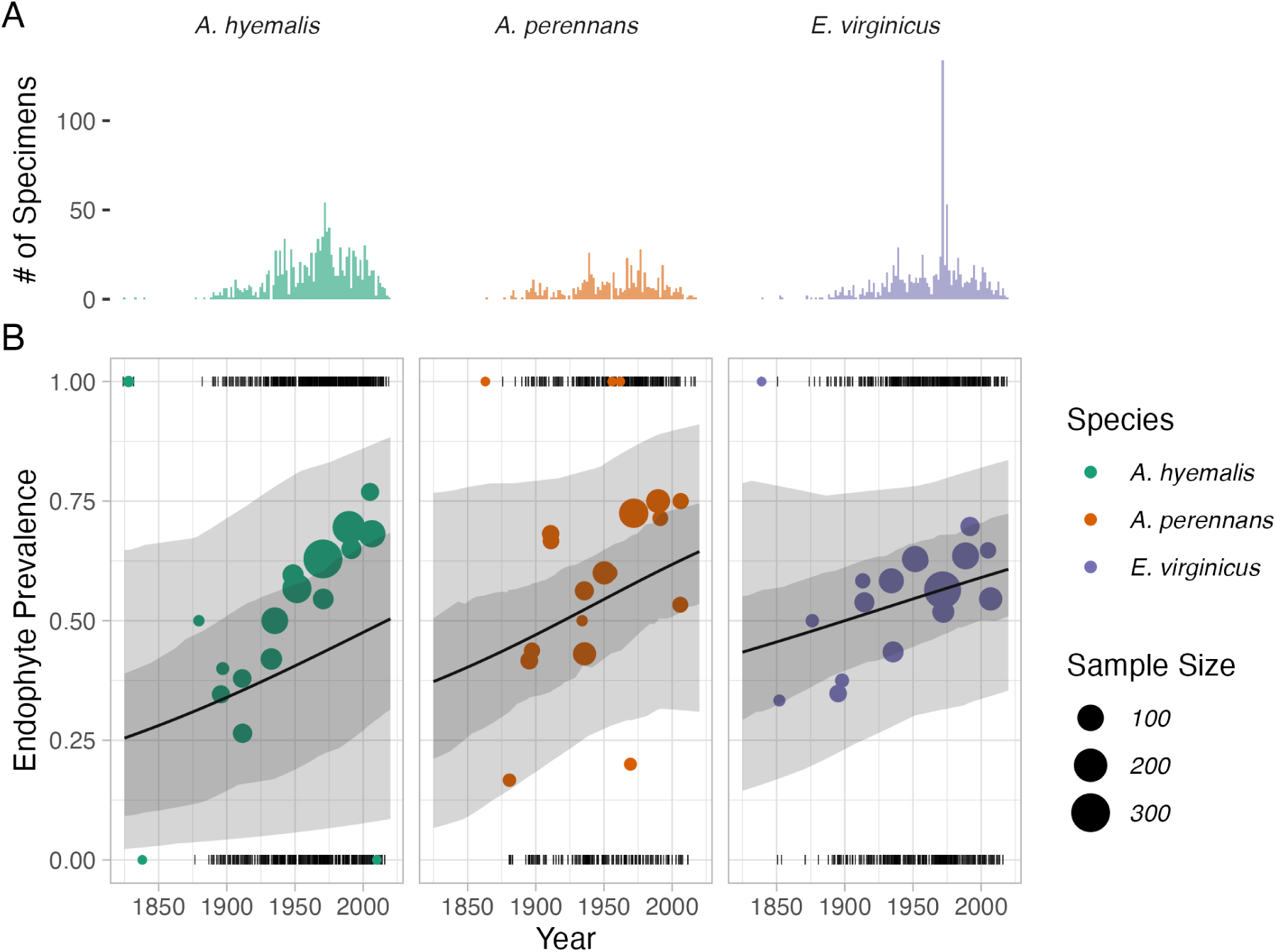
Temporal trends in endophyte prevalence. (A) Histograms show the frequency of scored specimens through time for each host species. (B) Lines show mean endophyte prevalence predicted by the endophyte prevalence model over the study period along with the 50% and 95% CI bands incorporating parameter uncertainty and variation associated with collector and scorer random effects. Colored points are binned means of the observed endophyte presence/absence data (black dashes). Colors represent each host species (*A. hyemalis*: green, *A. perennans*: orange, *E. virginicus*: purple) and point size represents the number of specimens.

Each specimen was assigned geographic coordinates based on collection information recorded on the herbarium sheet using the geocoding functionality of the *ggmap* R package (Kahle and Wickham, 2019). Many specimens had digitized collection information readily available, but for those that did not, we transcribed information printed on the herbarium sheet. The identity of each specimen collector was gathered as part of the sample’s metadata. Collections were geo-referenced to the nearest county centroid, or nearest municipality when that information was available. For fifteen of the oldest specimens, only information at the state level was available, and so we used the state centroid. The median pairwise distance between georeferenced coordinate points was 841 km. The median longitudinal width of the bounding boxes generated to geocode municipality, county, or state centroids was 44.7 km. Among those specimens geo-referenced at the state level, the largest bounding box, spanning the state of Texas, was 1233 km wide. The smallest bounding boxes were less than 1 km across for small municipalities (while this suggests high precision, we note that some specimens were collected in natural habitat nearby to small municipalities not encompassed by these bounding boxes).

Our visits focused on herbaria with historic strengths in grass collections (e.g. Texas A&M, Missouri Botanic Garden) and other herbaria in the Southern Great Plains region of the United States. While these nine herbaria garnered specimens that span the focal species’ ranges, our dataset unevenly samples across the study region (Fig. 1). Texas, Oklahoma, Louisiana, and Missouri are the most represented states. Uneven sampling was most pronounced for *A. perennans*, which has much of its range in the northeastern US. We explore the potential influence of spatial bias in sampling on our results through a simulation analysis (Appendix A - Supporting Methods).

After collecting seed samples, we quantified the presence or absence of *Epichloë* fungal hyphae in each specimen using microscopy. We first softened seeds with a 10% NaOH solution, then stained the seeds with aniline blue-lactic acid stain and squashed them under a microscope cover slip. We examined the squashed seeds for the presence of fungal hyphae at 200-400X magnification (Bacon and White, 2018). On average we scored 4.7 intact seeds per specimen of *A. hyemalis*, 4.2 seeds per specimen of *A. perennans*, and 3.8 seeds per specimen of *E. virginicus*; we scored 10, 342 seeds in total. Due to imperfect vertical transmission, the production of symbiont-free offspring from symbiotic hosts (Afkhami and Rudgers, 2008), it is possible that symbiotic host-plants produce a mixture of symbiotic and non-symbiotic seeds. We therefore designated a specimen as endophyte-symbiotic if *Epichloë* hyphae were observed in one or more of its seeds, or non-symbiotic if *Epichloë* hyphae were observed in none of its seeds. To capture uncertainty in the endophyte identification process, we recorded both a “liberal” and a “conservative” endophyte score for each plant specimen. When we confidently identified endophytes within a specimen’s seeds, we assigned matching liberal and conservative scores. When we identified potential endophytes with unusual morphology, low uptake of stain, or a small amount of fungal hyphae across the scored seeds, we recorded a positive identification for the liberal score and a negative identification for the conservative score. We recorded the identity of ech scorer as part of the data collection process. 89% of scored plants had matching liberal and conservative scores, reflecting high confidence in endophyte status. The following analyses used the liberal status, however repeating all analyses with the conservative status yielded qualitatively similar results (Fig. A8).

### Modeling spatial and temporal changes in endophyte prevalence

We assessed spatial and temporal changes in endophyte prevalence across each host distribution, quantifying the “global” temporal trends averaged across space, and then examining spatial heterogeneity in the direction and magnitude of endophyte change (hotspots and coldspots) across the spatial extent of each host’s distribution. To account for the spatial non-independence of geo-referenced occurrences, we used an approximate Bayesian method, Integrated Nested Laplace Approximation (INLA), to construct spatio-temporal models of endophyte prevalence. INLA provides a computationally efficient method of ascertaining parameter posterior distributions for certain models that can be formulated as latent Gaussian Models (Rue et al., 2009). Many common statistical models, including structured and unstructured mixed-effects models, can be represented as latent Gaussian Models. We incorporated spatial heterogeneity into this analysis using spatially-structured intercept and slope parameters implemented as stochastic partial differential equations (SPDE) to approximate a continuous spatial Gaussian process. This SPDE approach is a flexible method of smoothing across space while explicitly accounting for spatial dependence between data-points (Bakka et al., 2018; Lindgren et al., 2011). Fitting models with structured spatial effects is possible with MCMC sampling but can require long computation times, making INLA an effective alternative. This approach has been used to model spatial patterns in flowering phenology (Willems et al., 2022), the abundance of birds (Meehan et al., 2019) and butterflies (Crossley et al., 2022), the distribution of temperate trees (Engel et al., 2022) as well as the population dynamics of endangered amphibians (Knapp et al., 2016) and other ecological processes (Beguin et al., 2012).

We estimated global and spatially-varying trends in endophyte prevalence using a joint-likelihood model. For each host species *h*, endophyte presence/absence of the *i^th^* specimen (*P_h,i_*) was modeled as a Bernoulli response variable with expected probability of endophyte occurrence *P̂_h,i_*. We modeled *P̂_h,i_* as a linear function of collection year, with intercept *A_h_* and slope *T_h_* defining the global temporal trend in endophyte prevalence specific to each host species as well as with spatiallyvarying intercepts *α_h,l__i_* and slopes *τ_h,l__i_* associated with location (*l_i_*, the unique latitude-longitude combination of the *i*th observation). The joint-model structure allowed us to “borrow information” across species in the estimation of shared variance terms for the spatially-dependent random effect *δ_l__i_* , intended to account for residual spatial variation, and *χ_c__i_* and *ω_s__i_* , the i.i.d.-random effects indexed for each collector identity (*c_i_*) and scorer identity (*s_i_*) of the *i^th^* specimen.

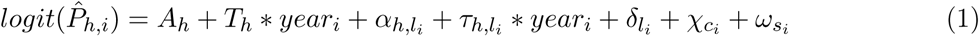

By including random effects for collectors and scorers, we accounted for “nuisance” variance that may bias predictions for changes in endophyte prevalence. Previous work suggests that behavior of historical botanists may introduce biases into ecological inferences made from historic collections (Kozlov et al., 2020). Prolific collectors who contribute thousands of specimens may be more or less likely to collect certain species, or specimens with certain traits (Daru et al., 2018). Similarly, the process of scoring seeds for hyphae involved multiple researchers (or “scorers”) who, even with standardized training, may vary in their likelihood of positively identifying *Epichloë*.

We performed model fitting using the *inlabru* R package (Bachl et al., 2019). Global intercept and slope parameters, *A* and *T* , were given vague priors. Collector and scorer random effects, *χ* and *ω* respectively, were centered at 0 with precision parameters assigned penalized complexity (PC) priors with parameter values *U_P C_* = 1 and *a_P C_* = 0.01 (Simpson et al., 2017). Each spatially-structured parameter depended on a covariance matrix according to the proximity of each pair of collection locations (Bakka et al., 2018; Lindgren et al., 2011). The covariance matrix was approximated using a Matérn covariance function, with each data point assigned a location according to the nodes of a mesh of non-overlapping triangles encompassing the study area (Fig. A2). Matérn covariance functions are widely used in spatially explicit statistical modeling because of their mathematical tractability and flexibility. This covariance structure relies on the assumption that the underlying process is stationary and isotropic, such that spatial autocorrelation between data points depends only on their relative positions (Bakka et al., 2018).

Implementing spatially-structured parameters in INLA with this SPDE approach is useful particularly because space is treated as a continuous variable, allowing the model to make efficient use of the data and generate predictions across the entire study region. The SPDE approach is flexible enough that it can capture smooth trends across space that are informed by the data rather than by spatial regions chosen *a priori* by researchers. However this flexibility also invites the risk of overfitting, as with other non-linear modeling approaches (Lapeyrolerie and Boettiger, 2023; Ramampiandra et al., 2023; Ward et al., 2014). Priors for the Matérn covariance function, termed “range” and “variance”, define how proximity effects decay with distance. The choice of priors for these types of spatial models is an area of active research (Bakka et al., 2018; Simpson et al., 2017), but another advantage of the INLA approach is that its computational efficiency allows for prior sensitivity analyses. Results presented in the main text reflect a prior range of 342 kilometers (i.e. a 50% probability of estimating a range less than 342 kilometers). We tested a range of values (from 68 kilometers to 1714 kilometers) and meshes (presented in the Supporting Methods – *Mesh and Prior Sensitivity Analysis*), finding that while the magnitude and uncertainty of effects varied, model results were qualitatively similar, i.e. the same direction of effects across space. We assessed model fit with visual posterior predictive checks (Fig. A3) and measurements of AUC (Figs. A4-A5) (Gelman and Hill, 2006). Through results and discussion that follow, we refer to the model described in this section as the “endophyte prevalence model”.

### Modeling distributions of host species

The herbarium records did not encompass the entirety of each host species’ range. Therefore, we used additional data sources to model the geographic distribution of each host species, with two goals: (1) generate realistic maps on which we could project the predictions of the INLA model, and (2) use the geographic distributions to test for relationships between climate change drivers and trends in endophyte prevalence. We followed the ODMAP (overview, data, model, assessment, prediction) protocol (Crossley et al., 2022) (see Supporting Methods). In short, we used presence-only observations of each host species from Global Biodiversity Information Facility (GBIF) between 1990 to 2020 (713 occurrence records for *A. hyemalis* (GBIF.Org, 2025a), 656 occurrence records for *A. perennans* (GBIF.Org, 2025b), and 2338 occurrence records for *E. virginicus* (GBIF.Org, 2025c)). We fit maximum entropy (MaxEnt) models using the maxent function in the R package *dismo* (Hijmans et al., 2017) using the following seasonal climate predictors (1990-2020 climate normals): mean and standard deviation of spring, summer, and autumn temperature, and mean and standard deviation of spring, summer, and autumn cumulative precipitation.

We generated 10,000 pseudo-absences as background points, and split the occurrence data into 75% for model training and 25% for model testing. The performance of models was evaluated with AUC (Jiménez-Valverde, 2012). We found AUC values of 0.862, 0.838, 0.821 respectively for *Agrostis hyemalis*, *Agrostis perennans*, and *Elymus virginicus* indicating good model fit to data. We used the training sensitivity (true positive rate) and specificity (true negative rate) to set a threshold for transforming the continuous predicted probabilities into binary presence - absence host distribution maps on which we projected INLA predictions of endophyte prevalence (Liu et al., 2005).

### Assessing the role of climate drivers

We assessed how the magnitude of climate change may have driven changes in endophyte prevalence by assessing correlations between changes in climate and changes in endophyte prevalence predicted from our spatial model at evenly spaced pixels across the study area.

We first downloaded monthly temperature and precipitation rasters from the PRISM climate group (Daly and Bryant, 2013) covering the time period between 1895 and 2020 using the *prism* R package (Hart and Bell, 2015). PRISM provides reconstructions of historic climate variables across the United States by spatially interpolating weather station data (Di Luzio et al., 2008). Because the magnitude of observed climate change differs across seasons, and because different growing seasons is a key feature of the biology of our focal host species, we calculated 30-year climate normals for seasonal mean temperature and cumulative precipitation for the recent (1990 to 2020) and historic (1895 to 1925) periods. We used three four-month seasons within the year (Spring: January, February, March, April; Summer: May; June, July, August; Autumn: September, October, November, December). This division of seasons allowed us to quantify differences in the primary climate change drivers, temperature and precipitation, associated with the two “cool” seasons, when we expected our focal species to be most active (*A. hyemalis* flowering phenology: spring; *E. virginicus*: spring and summer; *A. perennans*: autumn). In addition to mean climate conditions, environmental variability itself can influence population dynamics (Tuljapurkar, 1982) and changes in variability are a key prediction of climate change models (IPCC, 2021; Stocker et al., 2013). Therefore, we calculated the standard deviation for each annual and seasonal climate driver across each 30-year period. We then took the difference between recent and historic periods for the mean and standard deviation for each climate driver (Figs. A13-A15). All together, we assessed twelve potential climate drivers: the mean and standard deviation of spring, summer, and autumn temperature, as well as the mean and standard deviation of spring, summer, and autumn cumulative precipitation (the same climate covariates used in the MaxEnt models).

We then evaluated whether areas that have experienced the greatest changes in endophyte prevalence (hotspots of endophyte change) are associated with high degrees of change in climate (hotspots of climate change). To do so, we modeled the fitted, spatially-varying slopes of endophyte change through time (*τ_h,l_*) as a linear function of environmental covariates, with a Gaussian error distribution for a set of pixels across each host distribution. The continuous SPDE approach taken from our endophyte prevalence model allows us to generate predictions of temporal trends in prevalence at arbitrarily many pixels across each host distribution. Balancing computation time with resolution, we generated predicted trends for 546, 645, and 753 pixels across each host distribution for *A. perennans*, *A. hyemalis*, and *E. virginicus* respectively (pixel dimensions: *A. perennans* = 65 km x 36 km; *A. hyemalis* = 61km x 45 km; *E. virginicus* = 62 km x 40 km ). Fitting regressions to many pixels across the study region risks artificially inflating confidence in our results due to large sample sizes, and so we performed this analysis using only a random subsample of 250 pixels across the study region; other sizes of subsample yielded similar results. Data from each host species were analyzed separately. Throughout the results and discussion that follow, we refer to this analysis as the “*post hoc* climate regression analysis”.

### Validating model performance with in-sample and out-of-sample tests

We evaluated the predictive ability of the endophyte prevalence model using both in-sample training data from the herbarium surveys, and with out-of-sample test data, an important but rarely used strategy in ecological studies (Lee et al., 2024; Tredennick et al., 2021). We generated out-of-sample test data from contemporary surveys of endophyte prevalence in natural populations of *A. hyemalis* and *E. virginicus* in Texas and the southern US. Surveys of *E. virginicus* were conducted in 2013 as described in Sneck et al. (2017), and surveys of *A. hyemalis* took place between 2015 and 2020. Population surveys of *A. hyemalis* were initially designed to cover longitudinal variation in endophyte prevalence towards its range edge, while surveys of *E. virginicus* were designed to cover latitudinal variation. In total, we visited 43 populations of *A. hyemalis* and 20 populations of *E. virginicus* across the south-central US, with emphasis on Texas and neighboring states (Fig A12). number of plants sampled per population: 22.9); note that this sampling design provided greater local depth of information than the herbarium records, where only one plant was sampled at each locality. We quantified the endophyte status of each individual with microscopy as described for the herbarium surveys (with 5-10 seeds scored per individual), and calculated the prevalence of endophytes within the population (proportion of plants that were endophyte-symbiotic). For each population, we compared the observed fraction of endophyte-symbiotic hosts to the predicted probability of endophyte occurrence *P̂* derived from the model for that location and year. The contemporary survey period (2013-2020) is at the most recent edge of the time period encompassed by the historical specimens used for model fitting.

## Results

### How has endophyte prevalence changed over time?

Across more than 2300 herbarium specimens dating back to 1824, we found that prevalence of *Epichloë* endophytes increased over the last two centuries for all three grass host species (Fig. 2). On average, endophytes of *A. perennans* and *E. virginicus* increased from ∼ 40 % to 70% prevalence across the study region, and *A. hyemalis* increased from ∼ 25% to over 50% prevalence. Our model indicates high confidence that overall temporal trends are positive across species (99% probability of a positive overall year slope in *A. hyemalis*, 92% probability of a positive overall year slope in *A. perennans*, and 91% probability of a positive overall year slope in *E. virginicus*) (Fig. A6).

The model appears to under-predict the observed increase in endophyte prevalence relative to the data, particularly for *A. hyemalis* (Fig. 2B), but the model is accounting for random effects and spatial non-independence that are not readily seen in the figure. We found no evidence that collector biases influenced our results. Collector random effects were consistently small (Fig. A9), and models fit with and without this random effect provide qualitatively similar results. The identity of individual scorers, the researchers who identified endophyte status microscopically, did contribute to observed patterns in endophyte prevalence. For example, 3 of the 25 scorers were significantly more likely than average to assign positive endophyte status, as indicated by 95% credible intervals greater than zero, while 4 of the 25 had 95% credible intervals below zero (Fig. A10).

### How spatially variable are temporal trends in endophyte prevalence?

While there was an overall increase in endophyte prevalence, our model revealed hotspots and coldspots of change across the host species’ ranges, which are mapped in Fig. 3 across geographic ranges predicted by MaxEnt species distribution models. In some regions, posterior mean estimates of spatially varying temporal trends indicate that *A. hyemalis* and *A. perennans* experienced increases in prevalence by as much as 2% per year over the study period. Posterior estimates of uncertainty in spatially varying slopes indicate that these hotspots of change may have experienced increases of up to 5% per year while declines in prevalence may be as great as −4% per year for the *Agrostis* species. (Fig. A7) In contrast, *E. virginicus* experienced increases up to around 1% per year, with uncertainty ranging between 3.5% increases and 2.5% decreases (Fig. A7) Taken together, both *Agrostis* species show areas of both strong increasing and declining prevalence, while *E. virginicus* had an overall weaker and geographically more homogeneous increase in endophyte prevalence. Notably, endophytes are predicted to have increased most strongly towards the western range edge of *A. hyemalis* (Fig. 3A) and across the northeastern US for *A. perennans* (Fig. 3B). Broad increases in prevalence on average, along with increases towards range edges that had low historic prevalence result in range expansions of the symbiosis for both *Agrostis* species (Fig. 4). Increases in prevalence were strongest in regions with low historic prevalence for the *Agrostis* species (Fig. A11 A-B), but for *E. virginicus* trends did not differ according to historic prevalence (A11 C).

**Figure 3:**
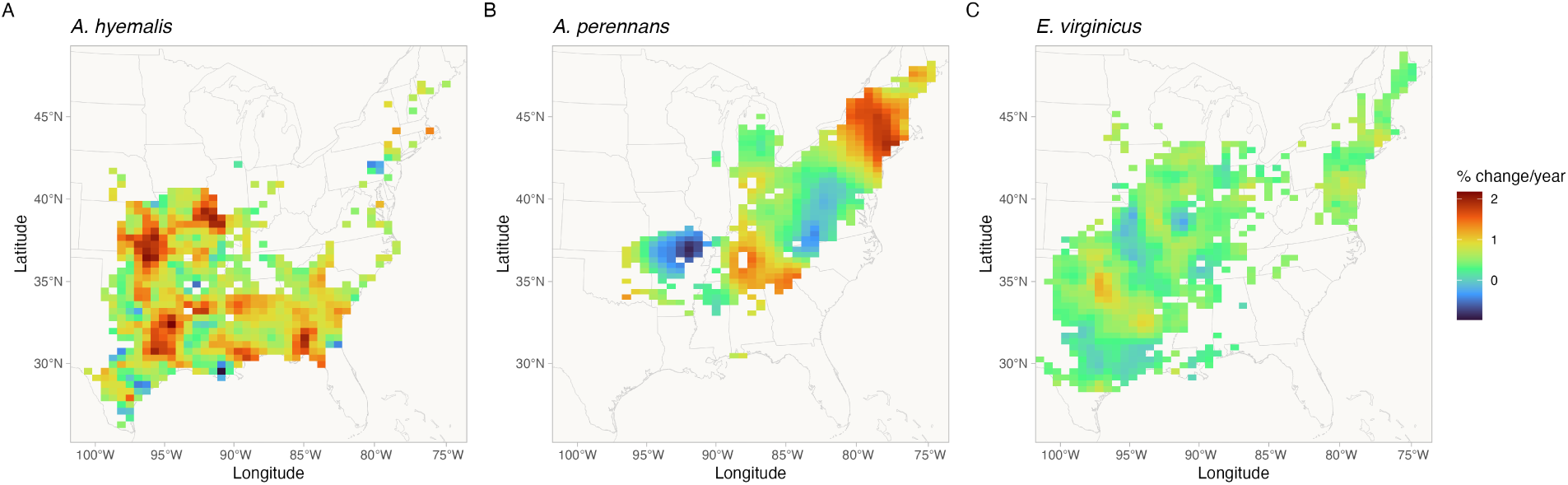
Predicted posterior mean of spatially-varying slopes representing change in endophyte prevalence for each host species ((A)*A. hyemalis*; (B)*A. perennans*; (C)*E. virginicus*). Spatially-varying trends are estimated from the endophyte prevalence model. Color indicates the relative change in predicted endophyte prevalence. Map lines delineate study areas and do not necessarily depict accepted national boundaries.

**Figure 4:**
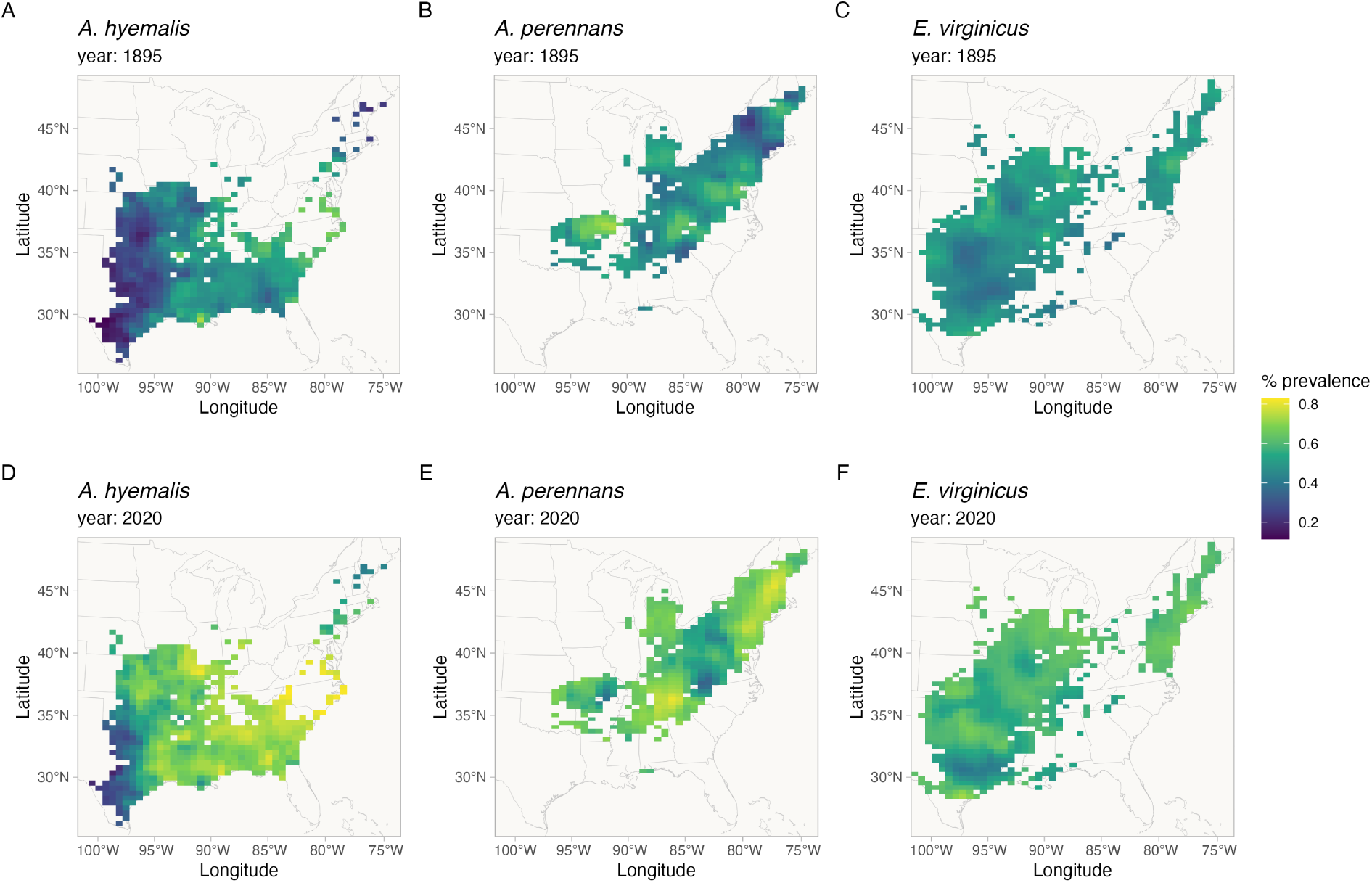
Predicted endophyte prevalence for each host species in 1895 and 2020. Predictions of prevalence come from the endophyte prevalence model. Color indicates the posterior mean endophyte prevalence for (A, D) *A. hyemalis*, (B, E) *A. perennans*, and (C, F) *E. virginicus*. Map lines delineate study areas and do not necessarily depict accepted national boundaries.

### What is the relationship between variation in temporal trends in endophyte prevalence and changes in climate drivers?

We found that trends in endophyte prevalence were strongly associated with one or more seasonal climate change drivers (Fig. 5). For the majority of the study region, the climate has become wetter (an average increase in annual precipitation of 60 mm) with relatively minimal temperature warming (an average increase in annual temperature of 0.02 ^◦^C) over the last century (Fig. A13-A15), a consequence of regional variation in global climate change (IPCC, 2021). Within the region, climate changes were spatially variable; certain locations experienced increases in annual precipitation as large as 375 mm or decreases up to 54 mm across the last century, while annual temperature changes ranged from warming as great as 1.4 ^◦^C to cooling by 0.46 ^◦^C.

**Figure 5:**
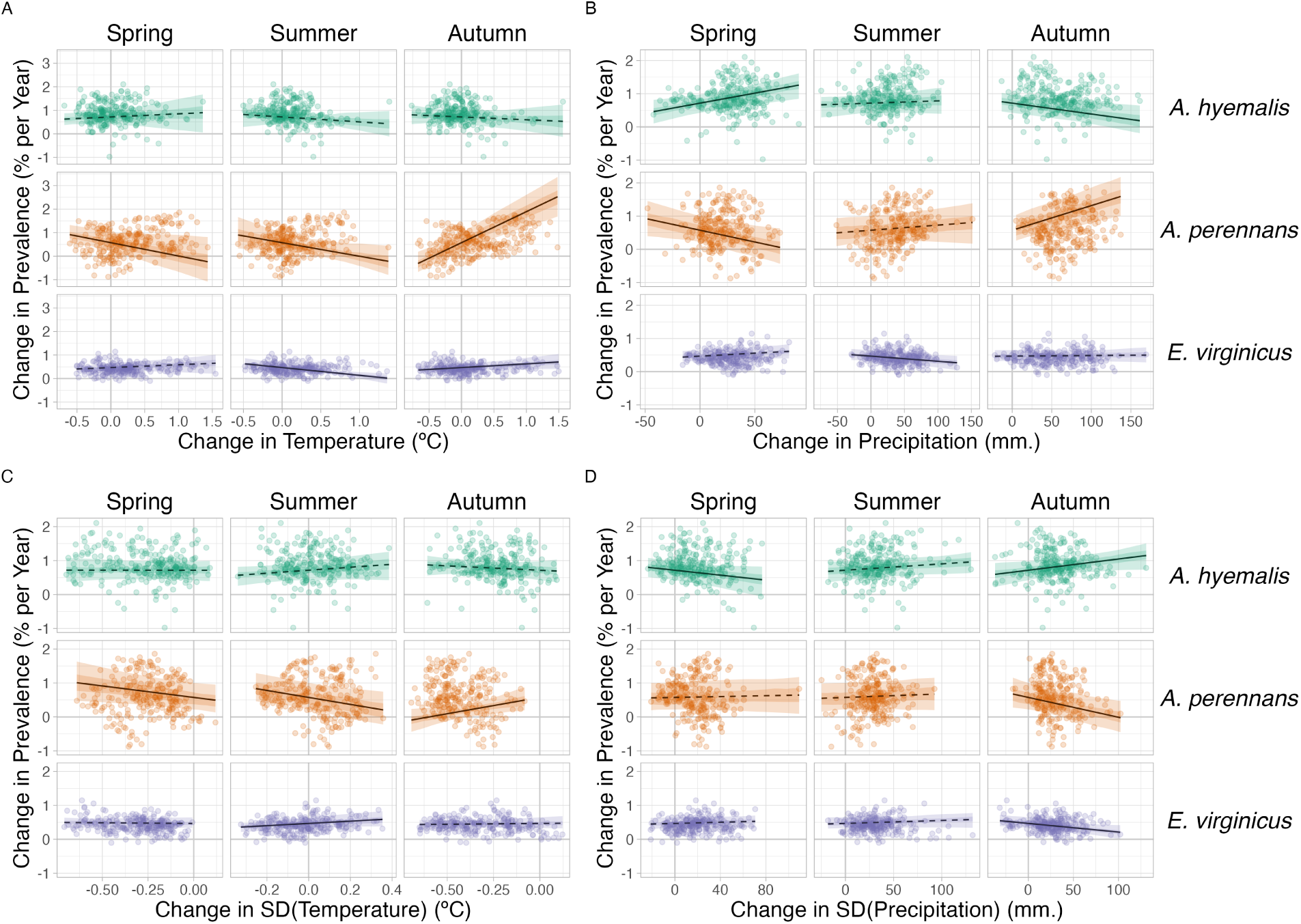
Relationships between predicted trends in endophyte prevalence and changes in seasonal climate drivers. Lines show marginal predicted relationship between spatially-varying trends in endophyte prevalence and changes in mean and variability of climate ((A): mean temperature, (B): cumulative precipitation, (C): standard deviation in temperature, (D): standard deviation in precipitation) estimated from the *post hoc* climate regression analysis. Confidence bands represent the 50 and 95% CI, colored by host species (*A. hyemalis*: green, *A. perennans*: orange, *E. virginicus*: purple). Slopes with greater than 95% posterior probability of being either positive or negative are represented as solid lines while those that have less than 95% probability are dashed. Points are the values of pre-computed SVC trends and climate drivers at 250 randomly sampled pixels across each host’s distribution used in model fitting for the *post hoc* climate regression analysis.

Spatially variable climate trends were predictive of trends in endophyte prevalence. For example, among the tested climate drivers, strong increases in endophyte prevalence for *A. perennans* were most strongly associated with increasing autumn precipitation and with increasing mean and variability in autumn temperature (greater than 97% posterior probabilities of positive slopes). For this species, each 1 ^◦^C increase in autumn temperature was associated with a 1.07 % greater increase per year in endophyte prevalence (Fig. 5A) and a 100 mm increase in precipitation was associated with a 0.8% greater increase per year in endophyte prevalence (Fig. 5B). This result aligns with the species’ autumn active growing season, however other seasonal climate drivers were also positively associated with increasing endophyte prevalence in this host species. In particular, we found cooler and drier springs and cooler summers to be associated with increasing endophyte prevalence (greater than 99% posterior probabilities of negative slopes), though these slopes were generally of smaller magnitude than those for autumn climate drivers. Changes in endophyte prevalence across the ranges of *A. hyemalis* and *E. virginicus* were less strongly driven by changes in climate. Like *A. perennans*, climate during peak growing season (spring for *A. perennans* and summer for *E. virginicus*) emerged most commonly as drivers of changes in endophyte prevalence. Across the tested climate drivers, increases in mean spring precipitation were the strongest predictor of increasing trends in endophyte prevalence for *A. hyemalis* (Fig. 5B) (greater than 99% posterior probability of a positive slope). For this species, an increase of 100 mm in spring precipitation was associated with 0.6% per year stronger increases in endophyte prevalence relative to regions with no change in precipitation. The next greatest slopes were those associated with variability in spring precipitation (greater than 96% posterior probability of a negative slope), as well as in the mean and variability of autumn climate (greater than 98% probability of negative and positive slopes, respectively).

Changes in endophyte prevalence in *E. virginicus* were not strongly associated with changes in most climate drivers, but regions with reduced variability in autumn precipitation (Fig. 5B) and with cooler and more variable summer temperatures (Fig. 5A,C) experienced the largest increases in endophyte prevalence. Our analysis indicated relatively high confidence that these climate drivers influence endophyte prevalence shifts in *E. virginicus*(greater than 99% posterior probability of either negative or positive slopes respectively), however they translate, for example, to less than a 0.4% decrease in endophyte prevalence per year for each 1*^o^C* of summer warming over the century. Repeating this analysis using all pixels across each species’ distribution were qualitatively similar to these results.

### Evaluation of model performance on an out-of-sample test

Tests of the endophyte prevalence model’s predictive performance, as quantified by AUC and by visual posterior predictive checks, indicated good predictive ability. Model performance was similar between historic herbarium specimens used as training data and the out-of-sample test data from contemporary surveys (AUC = 0.79 and 0.77 respectively; Fig. A5-A4). The model successfully captured broad regional trends in endophyte prevalence seen in the contemporary survey data, such as decline endophyte prevalence in *A. hyemalis* towards western longitudes (Fig. 6A) and an increase towards northern latitudes (Fig. 6B). It is noteable that model predictions for endophyte prevalence exhibited relatively little local geographic variation, whereas the out-of-sample survey data were highly variable with populations spanning 0% to 100% endophyte-symbiotic plants (Fig. 6C), indicating population-to-population variation not captured in the endophyte prevalence model.

**Figure 6:**
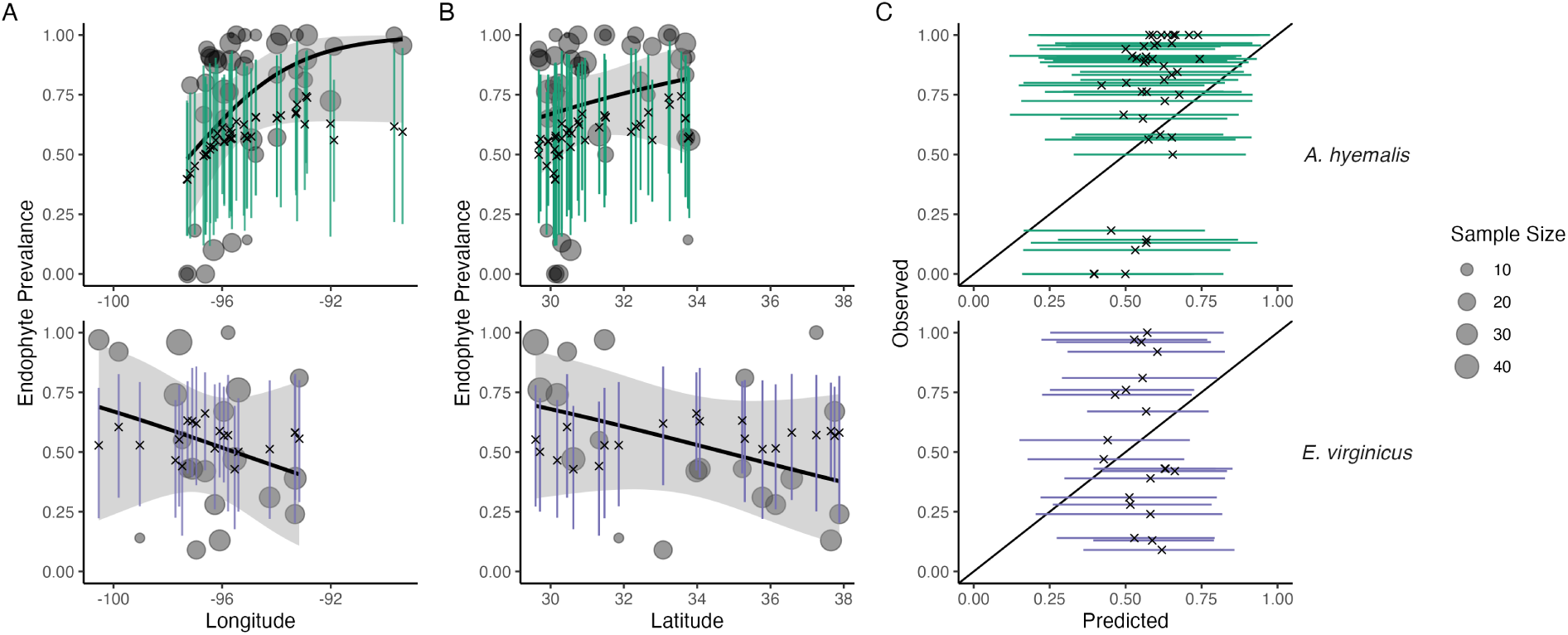
Predictive performance for contemporary test data. (A) The endophyte prevalence model, trained on historic herbarium collection data, performed modestly at predicting prevalence in contemporary population surveys. The model captured regional trends across (A) longitude and (B) latitude. Crosses indicate predicted mean prevalence along with the 95% CI (colored lines: *A. hyemalis*: green, orange, *E. virginicus*: purple) from the herbarium model. Contemporary prevalence is represented by grey points (point size reflects sample size) along with trend lines from generalized linear models (black line and shaded 95% confidence interval). (C) Comparison of contemporary observed population prevalence vs. predicted endophyte prevalence shows that contemporary test data had more variance between populations than in model predictions.

## Discussion

Our examination of historic plant specimens revealed previously hidden shifts in microbial symbiosis over the last two centuries. For the three grass host species we examined, there have been strong increases in prevalence of *Epichloë* endophyte symbiosis. We interpret increases in prevalence of *Epichloë*, which are vertically transmitted, as adaptive changes that improve the fitness of their hosts under increasing environmental stress. This interpretation is in line with theory predicting that positive fitness feedback caused by vertical transmission leads beneficial symbionts to rise in prevalence within a population (Donald et al., 2021; Fine, 1975). We further found that trends in endophyte prevalence often varied across the host distribution in association with changes in climate drivers, consistent with the hypothesis that increases in endophyte prevalence are driven by context-dependent benefits to hosts that confer resilience under environmental change. Taken together, our results suggest an overall strengthening of host-symbiont mutualism over the last two centuries.

### Responses of host-microbe symbioses to climate change

Differences across host species underscore that while all of these *C*_3_ grasses share similar broad-scale distributions, each engages in unique biotic interactions and has unique responses to environmental drivers. We identified hotspots of change for *A. perennans*, which was the species whose endophyte prevalence was most responsive to changes in climate drivers (Fig. 5). Predicted declines of 0.9% per year in the southern portion of its range and predicted increases of up to 2% per year in the north suggest a potential poleward range shift of endophyte-symbiotic plants (Fig. 3B); whether the overall host distribution is shifting in parallel is an exciting next question.

Based on previous work demonstrating that endophytes can shield their hosts from drought stress (reviewed in Decunta et al. (2021)), we generally predicted that drought conditions would be a driver of increasing endophyte prevalence. In contrast to this expectation, increasing prevalence for *A. perennans* was associated with both increasing autumn temperature and precipitation (Fig. 5). To our knowledge, the response of the symbiosis in *A. perennans* to drought has not been examined experimentally, but in a greenhouse experiment, endophytes had a positive effect on host reproduction under shaded, low-light conditions (Davitt et al., 2010). Our results also hint that it may be useful to investigate whether lagged climate effects are important predictors of host fitness in this system (Evers et al., 2021). Endophyte prevalence of the autumn-flowering *A. perennans* was strongly linked with decreasing spring precipitation, and that of the spring-flowering *A. hyemalis* was associated with decreasing autumn precipitation (Fig. 5B). For *A. hyemalis*, endophytes could be playing a role helping hosts weather autumn-season droughts, which is likely also an important time for the species’ germination. Previous work demonstrated drought benefits in a greenhouse manipulation with this host-symbiont pair (Davitt et al., 2011), and early life stages may be particularly vulnerable to prolonged droughts. For *E. virginicus*, which experienced the weakest changes in endophyte prevalence overall (ranging between 1.1% increases and 0.2% decreases), we only found modest associations with changes in climate drivers. Surveys by Sneck et al. (2017), used as part of the test data in this study, identified a drought index (SPEI) that integrates precipitation with estimated evapotranspiration as an important predictor of contemporary endophyte prevalence in this species. The diverse relationships we detect between trends in endophyte prevalence and climate drivers suggest a more complicated picture than the simple explanation that drought alone, as measured through changes in annual precipitation, causes increasing endophyte prevalence through context-dependent fitness benefits.

While we show consistent increasing trends in prevalence between the three species, the mechanisms that explain these changes may be diverse and idiosyncratic. First, climate change responses may depend on genotype-specific responses that are not considered in our current analysis. While *Epichloë* symbioses are highly specialized, surveys have demonstrated genotypic and chemotypic diversity of the symbionts among and within populations (Treindl et al., 2023; von Cräutlein et al., 2021). Genotypic variation in *Epichloë* endophytes, particularly in genes responsible for alkaloid production, produces “chemotypes” with differing benefits for hosts against insect or mammalian herbivores mediated by environmental conditions (Saikkonen et al., 2013; Schardl et al., 2012). Genotypic variation of the hosts themselves can also influence interaction outcomes (Gundel et al., 2011; Parker et al., 2017). Whether hotspots of change in endophyte prevalence reflect selection for genotype-pairings with particularly strong fitness benefits is an unanswered question. Additionally, *Epichloë* endophytes have been connected to a suite of non-drought related fitness benefits including herbivory defense (Brem and Leuchtmann, 2001), salinity resistence (Wang et al., 2020), and mediation of pathogens (Vikuk et al., 2019) and the soil microbiome (Roberts and Ferraro, 2015). Broad changes in the distribution and abundance of natural enemies (Côté et al., 2004), along with stresses from anthropogenic changes in land management and pollution (Sage, 2020) likely influence the benefits of symbiosis (Rudgers et al., 2020). Changing endophyte prevalence results from the combination of net fitness benefits playing out across the heterogeneous map of a changing climate and and its interactive effects on other anthropogenic drivers. Host species experience a world that is made increasingly stressful by a combination of global change drivers, and while historic trends that we observed suggest that symbiotic fitness benefits have helped mitigate this stress, it is possible that at yet higher levels of stress, increasing costs of the mutualism could lead to declines in endophyte prevalence. More extreme climate stresses, which are expected more frequently in the future (Seneviratne et al., 2021) could shift the balance of interactions costs and benefits. Identifying ‘tipping points’ of mutualism breakdown under increasing environmental stress is an important area of future inquiry.

Our results indicate that *Epichloë* symbiosis has likely improved host fitness in stressful environments leading to increasing prevalence. What is less clear is how this will influence future range shifts. Based on our analysis, it is likely that the symbiosis will facilitate range shifts for hosts by improving population growth at range edges. Previous population surveys (Rudgers and Swafford, 2009; Semmartin et al., 2015; Sneck et al., 2017) attributed environment-dependent gradients in endophyte prevalence to symbiont-derived fitness benefit’s allowing hosts to persist in environments where they otherwise could not (Afkhami et al., 2014; Kazenel et al., 2015). However, symbiont-facilitated range shifts require that endophytes be present in the populations to be able to contribute to population growth. For example, the arid western range edge of *A. hyemalis* has had historically low endophyte prevalence (Fig. 4), and while prevalence has increased most quickly in regions with historically low endophyte prevalence (Fig. A11), the complete absence of endophytes at range edges in this species would make it impossible for prevalence to increase without dispersal of symbiotic seeds (Fowler et al., 2023). These factors potentially contribute to the ability of the host species to track its environmental niche. Another interesting question is the degree to which symbiotic and non-symbiotic hosts, which occupy overlapping but distinct niches, are likely to experience distribution shifts in tandem or at different rates in the future.

### Steps towards forecasts of host-microbe symbioses

The combination of a spatially-explicit model and historic herbarium specimens allowed us to identify regions of both increasing and decreasing endophyte prevalence. We see several next steps toward the goal of predicting host and symbiont niche-shifts in response to future climate change. While the model successfully predicted large-scale spatial trends observed in the outof-sample contemporary population surveys, these data contained more population-to-population variability in prevalence than could be explained by the model. We interpret this to mean that the model captures coarse-scale spatial and temporal trends reasonably well, but is not equipped to capture local-scale nuances that generate population-to-population differences. Validating our model predictions with this test, a rare extra step in collections-based studies, allows us to identify ways in which the model’s out-of-sample predictive ability could be improved. Lack of information on local variability in symbiont prevalence may simply be a feature of data derived from herbarium specimens. Natural history collectors sample one or a few specimens from local populations, and these observations are aggregated by the model to derive broad-scale estimates. This suggests that increasing local replication should be a factor considered in future collection efforts of natural history specimens, balancing the required time and effort along with limitations on storage space within collections. Herbarium collections were predominately used for taxonomic research in the past, but use of specimens to understand ongoing global change would benefit from increased collection efforts and expansion of herbarium collections. An alternative validation test would be to hold-out samples from the historic data set. Such a test would more clearly match the conditions of the training data (i.e., in spatial scale and climate conditions), however the trade-off between training and testing the model with a limited number of sampled specimens held us back from exploring this option. Splitting datasets can negatively impact model estimates, and the choice of how to split the data for model validation is not trivial (Bergmeir and Benítez, 2012; James et al., 2013).

Another key consideration in forecasting the dynamics of host-microbe symbioses is the spatial scale of both specimen georeferencing and available climate data. For this analysis, most specimen localities were assigned coordinates at county or city centroids, and the climate data examined was on 4 km grid cells. Georeferencing of specimens as accurately as possible is a key priority of herbarium specimen digitization efforts (Davis, 2023; Soltis, 2017). While the INLA modeling approach that we used allows for predictions at arbitrarility small spatial scales, and would simplify connecting model predictions to the scale of a given climate driver, the course scale inherent to our analysis may obscure some local-scale ecological processes Poor predictive ability at local scales in this grass-endophyte system is not surprising, as previous studies have found that local variation (e.g., in soil conditions, in microclimate), even to the scale of hundreds of meters can structure endophyte-host niches (Gundel et al., 2024; Kazenel et al., 2015). Local adaptation in either the host or symbiont to microclimate or soil conditions could cause populations to differ from broad regional trends. The choice of prior distributions for spatially-varying random effects also impacts the model’s flexibility to capture spatial trends. Our exploration of model sensitivity to prior choice(presented in the *Supplemental Methods* ) reveals qualitatively similar results across a broad range of priors. An important next step would be integrating data from local and regional scales through modeling to constrain estimates of local and regional variation.

Predicting future niche-shifts of hosts and symbionts will require considering the coupled dynamics of host-symbiont dispersal in addition to fitness benefits. For example, transplanting symbiotic and non-symbiotic plants beyond the range edge of *A. hyemalis* could tell us whether low endophyte prevalence in that area (Fig. 4A) is a result of environmental conditions that lead the symbiosis to have negative fitness consequences, or is a result of some historical contingency or dispersal limitation that has thus far limited the presence of symbiotic hosts from a region where they would otherwise flourish and provide resilience. Incorporating available climatic and soil layers as covariates is another obvious step that could improve predictions. These steps will bridge gaps that often exist between large but broad bioclimatic and biodiversity data and small but high-resolution data on biotic interactions, and move towards the goal of predicting the dynamics of microbial symbioses under climate change (Isaac et al., 2020; Miller et al., 2019).

### Herbaria for global change research

Our analysis advances the use of herbarium specimens in global change biology in two ways. First and foremost, this is one of a growing number of studies to examine microbial symbiosis using specimens from natural history collections, and the first, to our knowledge, to link long-term changes in symbioses to changes in climate. The responses of microbial symbioses are a rich target for future studies within historic specimens, particularly those that take advantage of advances in sequencing technology. While we used relatively coarse presence/absence data based on fungal morphology, other studies have examined historic plant microbiomes using molecular sequencing and sophisticated bioinformatics techniques, but these studies have so far been limited to relatively few specimens at limited spatial extents (Bearchell et al., 2005; Bieker et al., 2020; Bradshaw et al., 2021, 2023; Gross et al., 2021; Heberling and Burke, 2019; Yoshida et al., 2015). Much of this work highlights the important role that historic specimens can play in tracking pathogens, a particularly important area as climate change facilitates the spread of new diseases (Ristaino, 2020; Singh et al., 2023) Continued advances in capturing historic DNA and in filtering out potential contamination during specimen storage (Bakker et al., 2020; Daru et al., 2019; Raxworthy and Smith, 2021) will be imperative in the effort to scale up these efforts. This scaling up will be essential to be able to quantify changes not just in the prevalence of symbionts, but also in symbionts’ intraspecific variation and evolutionary responses to climate change, as well as in changes in the wider host microbiome. With improved molecular insights from historic specimens, we could ask whether the broad increases in endophytes that we have identified reflect selection for particular genetic strains or chemotypes and how this selection varies across space. Answering these questions as well as the unknown questions that future researchers may ask also reiterates the value in capturing meta-information during ongoing digitization efforts at herbaria around the world and during the accession of newly collected specimens (Edwards et al.; Lendemer et al., 2020).

The second major advance in this analysis is in accounting for several potential biases in the data observation process that may be common to many collections-based research questions by using a spatially-explicit random effects model. Potential biases introduced by the sampling habits of collectors (Daru et al., 2018), and variation between contemporary researchers during the collection of trait data, if not corrected for could lead to over-confident inference about the strength and direction of historic change (Fig. 2). Previous studies that have quantified the effects of collector biases typically find them to be small (Davis et al., 2015; Meineke et al., 2019), and we similarly did not find that collector has a strong effect on the results of our analysis, but that scorer identity did impact results. It is difficult to distinguish whether the impact of scorers was driven by true differences in scorers’ biases or by unintended spatial or temporal clustering of the specimens examined by each scorer (Clayton et al., 1993; Urdangarin et al., 2023). By under-weighting endophyte-positive samples that are clustered spatially or by collector or observer, the endophyte prevalence model is appropriately accounting for nuisance variables and providing a conservative inference of endophyte change relative to the raw data. Spatial autocorrelation is another phenomenon likely common in data derived from herbarium specimens (Willems et al., 2022), which our spatially-explicit analysis models among samples. Beyond spatial autocorrelation of outcomes, systematic differences in sampling across space can result in spatial bias.

One strength of herbaria as vehicles for global change research is the relative ease with which specimens from many distinct geographic locations can be examined. We visited just nine institutions in the central southern United States, and we were able to sample seeds from specimens across an area spanning over 300,000 sq. km, including specimens from Mexico and Canada. Despite this advantage, the specimens we examined are concentrated in the south-central United States, with fewer specimens in the rapidly warming northeastern United States reflecting the regional focus of herbaria. We provide a simulation analysis exploring the potential impact of spatially and temporally biased sampling (Appendix A - Supporting Methods). We found that the spatially-varying coefficient model had a strong ability to re-capitulate temporal trends across space in simulated data, and that this result was robust to relatively high levels of spatial bias (80% of data missing from one spatial region). Simulation analyses that extend this work to consider the myriad ways herbarium data may be biased (i.e. testing different spatial arrangements and scales of spatial bias, or testing different sample sizes) would be extremely valuable (Daru et al., 2018; Erickson and Smith, 2021; Gaul et al., 2020; Meineke and Daru, 2021; Schmidt et al.).

## Conclusion

Ultimately, a central goal of global change biology is to generate predictive insights into the future of natural systems on a rapidly changing planet. Beyond host-microbe symbioses, detecting ecological responses to anthropogenic global change and attributing their causes would inform public policy decision-makers and adaptive management strategies. Natural history specimens, such as the plant hosts examined in this study, have a clear role to play in informing global change biodiversity science, including building understanding of the dynamics of host-symbiont interactions (Davis, 2023). This survey of historic endophyte prevalence is necessarily correlative, yet it serves as a foundation to develop better predictive models of the response of microbial symbioses to climate change. Combining the insights from this type of regional-scale survey with field experiments and physiological performance data could be invaluable to identify mechanisms driving shifts in host-symbiont dynamics. Evidence is strong that certain dimensions of climate change correlated with endophytes’ temporal responses, however we do not know why trends in prevalence were weak in some areas or how endophytes would respond to more extreme changes in climate. The “time machine” of natural history collections revealed evidence of mutualism resilience for grass-endophyte symbioses in the face of environmental change, but more extreme changes could potentially push one or both partners beyond their physiological limits, leading to the collapse of the mutualism; more research is needed to understand what those limits might be.

## Acknowledgments

We thank Dr. Jessica Budke for help in drafting our initial destructive sampling plan, and to the many staff members of herbaria who facilitated our research visits, as well as to the hundreds of collectors who contributed to the natural history collections. Several high school and undergraduate researchers contributed to data collection, including A. Appio-Riley, P. Bilderback, E. Chong, K. Dickens, L. Dufresne, B. Gutierrez, A. Johnson, S. Linder, E. Scales, B. Scherick, K. Schrader, E. Segal , G. Singla, and M. Tucker. This research was supported by funding from National Science Foundation (grants 1754468 and 2208857) and by funding from the Texas Ecolab Program. Four anonymous reviewers greatly improved earlier versions of this manuscript. Statement of Authorship J.C.F. contributed to research conception, data collection, data analysis, and led manuscript drafting. J.M. contributed to data analysis and manuscript revisions. T.E.X.M. contributed to research conception, data collection, data analysis, and manuscript revisions.

## Data and Code Availability

Data from this publication can be found through a publicly available repository (https://doi.org/10.5061/dryad.rn8pk0pn0). Code for analyses can be found through a publicly available repository (https://github.com/joshuacfowler/EndoHerbarium) that will be permanently archived upon publication.

## Appendix A

### Supplemental Figures

**Figure A1:**
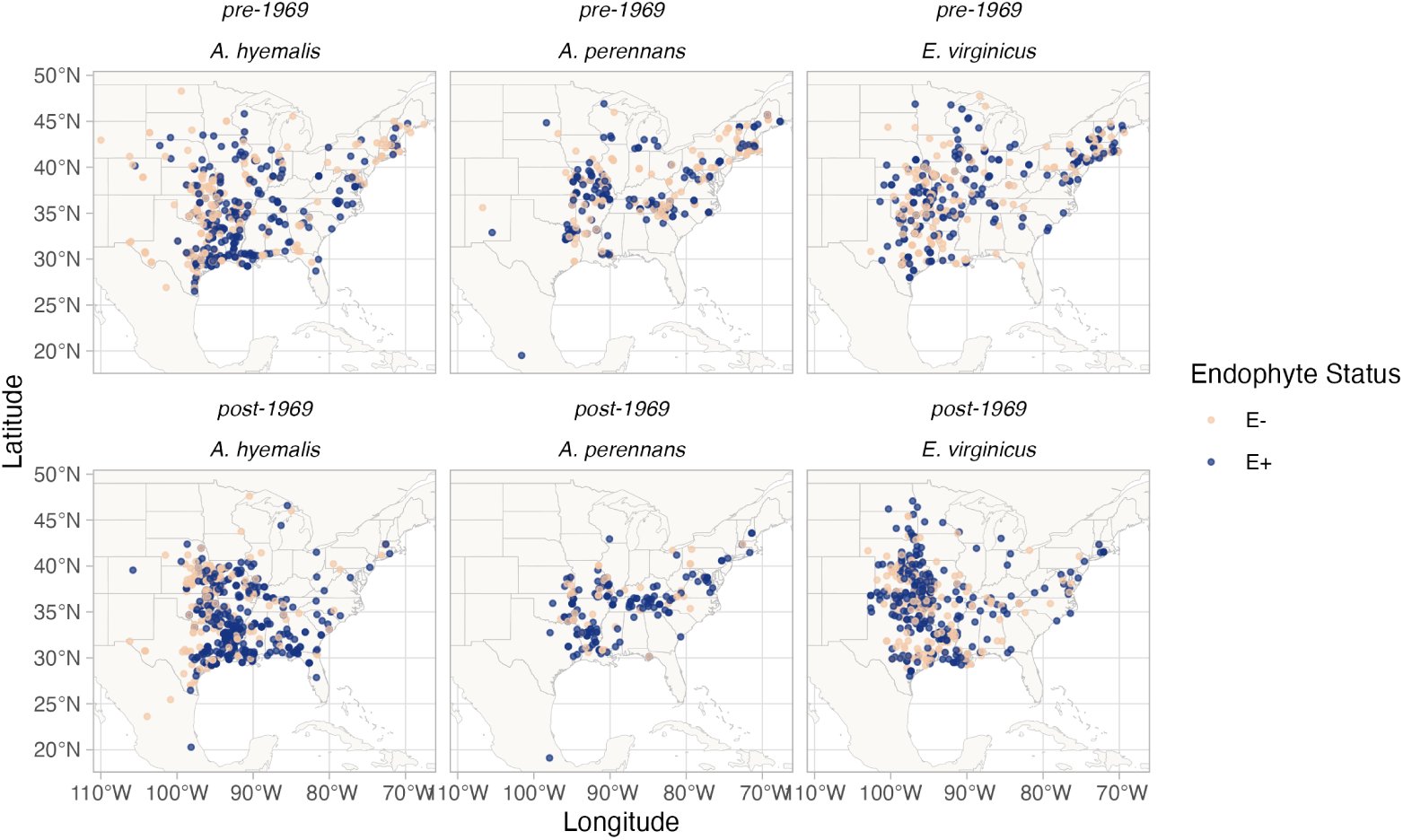
Endophyte presence/absence in specimens of each host species. Points show collection locations colored according to whether the specimen contained endophytes ( E+; blue points) or did not contain endophytes (E-, tan points). To visualize temporal change, the data are faceted before and after the median year of collection. Map lines delineate study areas and do not necessarily depict accepted national boundaries.

**Figure A2:**
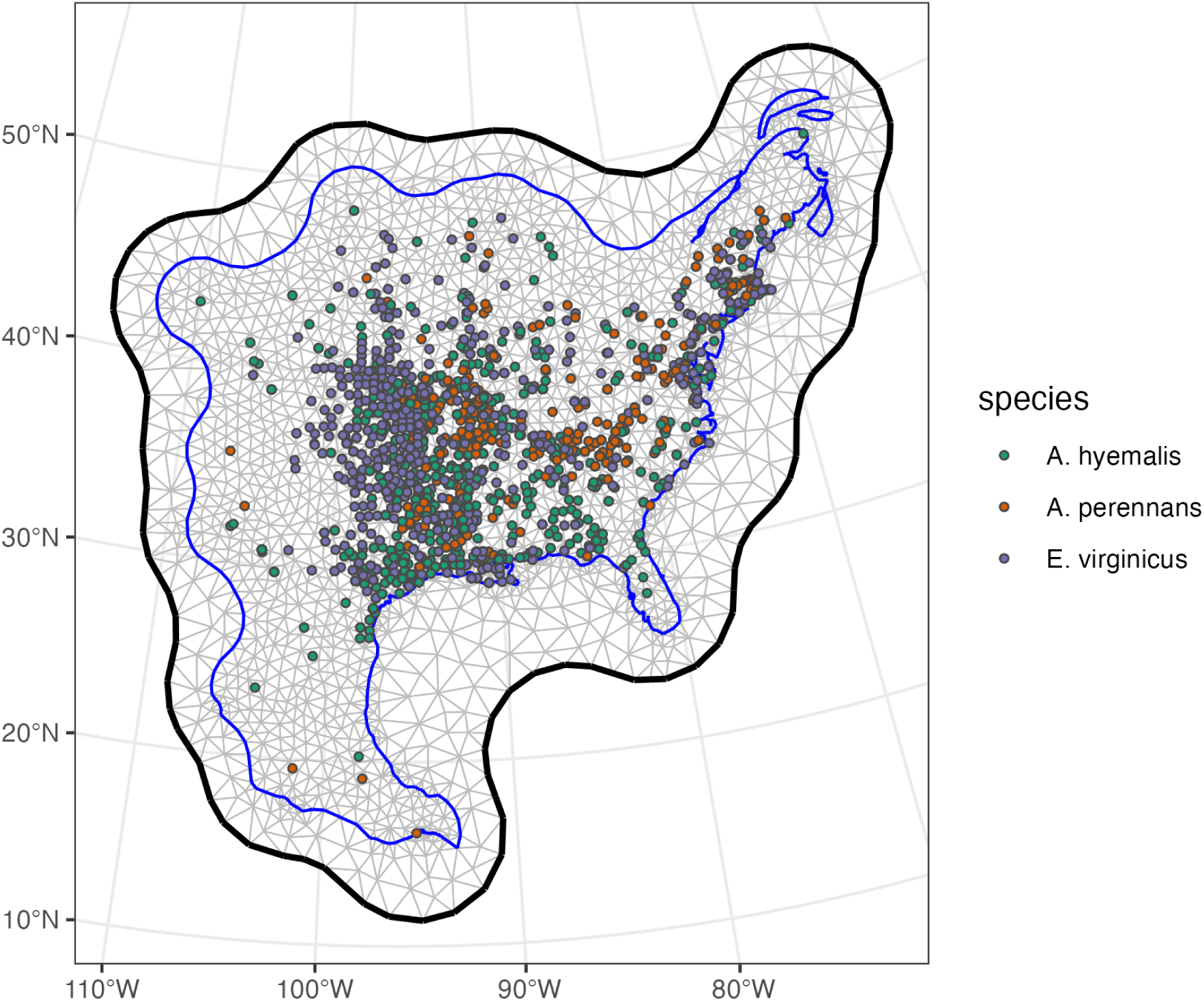
Triangulation mesh used to estimate spatial dependence between data points. Grey lines indicate edges of triangles used to define distances between observations. Colored points indicate locations of sampled herbarium specimens for each host species, and the blue line shows the convex hull and coastline used to define the edge of the mesh around the data points. The thick black line shows the convex hull defining a buffer space around the edge of the mesh to reduce the influence of edge effects on model estimates.

**Figure A3:**
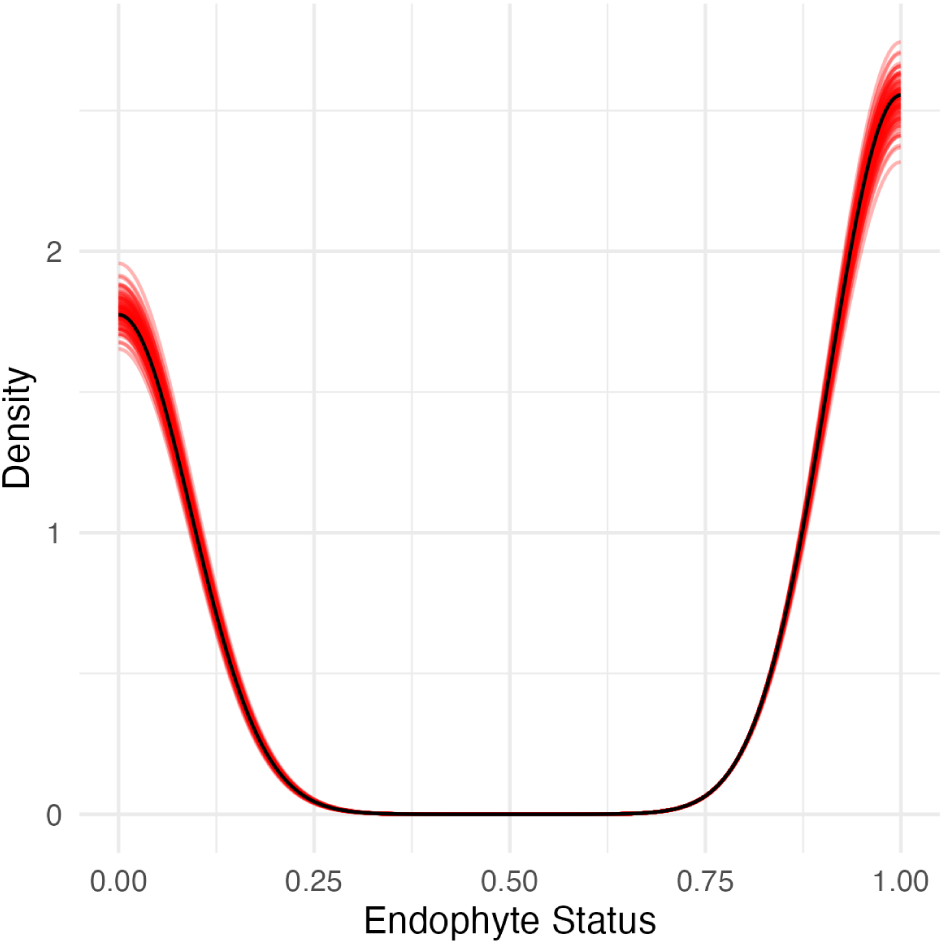
Graphical posterior predictive check of the endophyte prevalence model fit. Consistency between observed data and predicted values indicate that the fitted model accurately describes the data. Graph shows density curves for the observed data (black) along with 100 predicted datasets (red).

**Figure A4:**
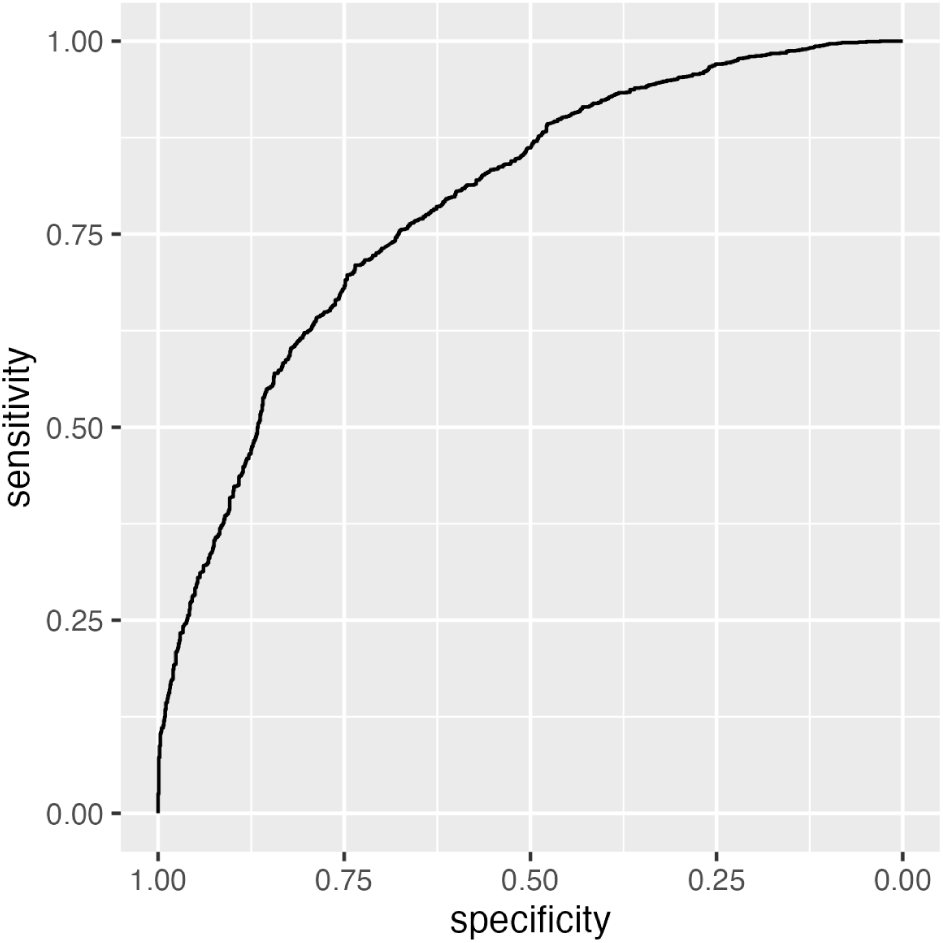
ROC plot showing performance of the endophyte prevalence model in classifying observations according to endophyte status within the in-sample training data from herbarium collections. The curves show adequate model performance for observed data. The AUC value is 0.79.

**Figure A5:**
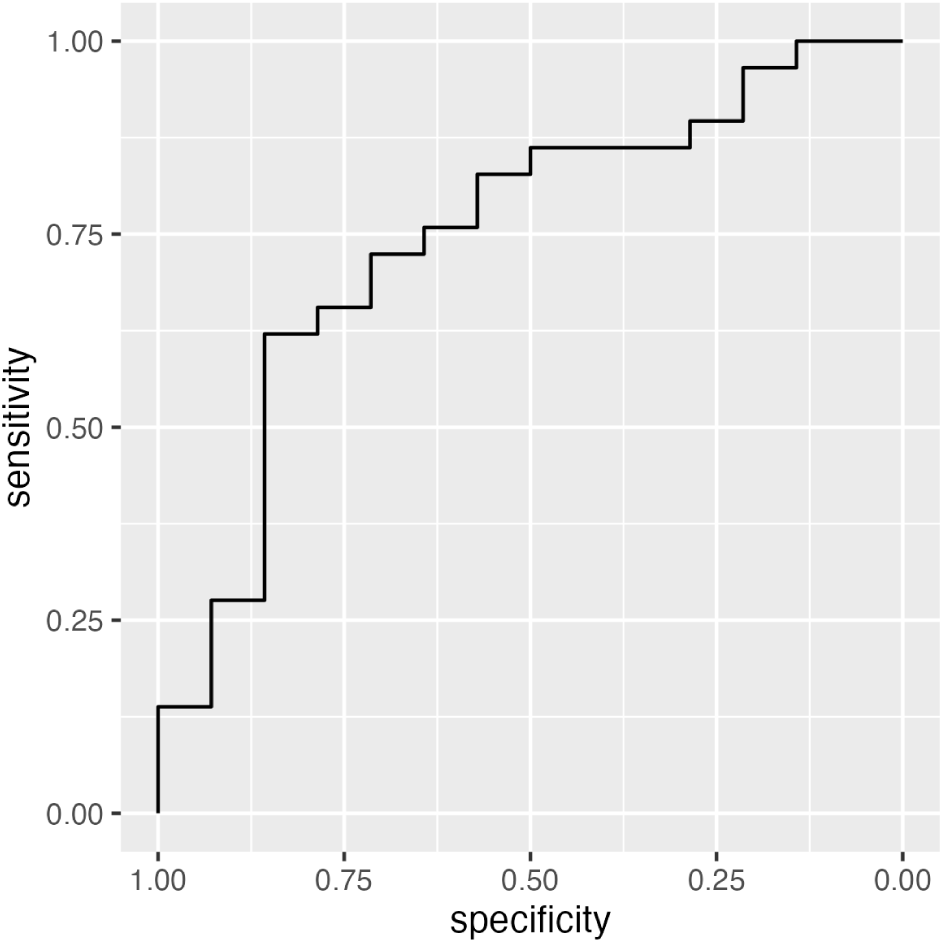
ROC plot showing performance of the endophyte prevalence model in classifying observations according to endophyte status within the out-of-sample test data from contemporary surveys. The curves show adequate model performance for test data. The AUC value is 0.77.

**Figure A6:**
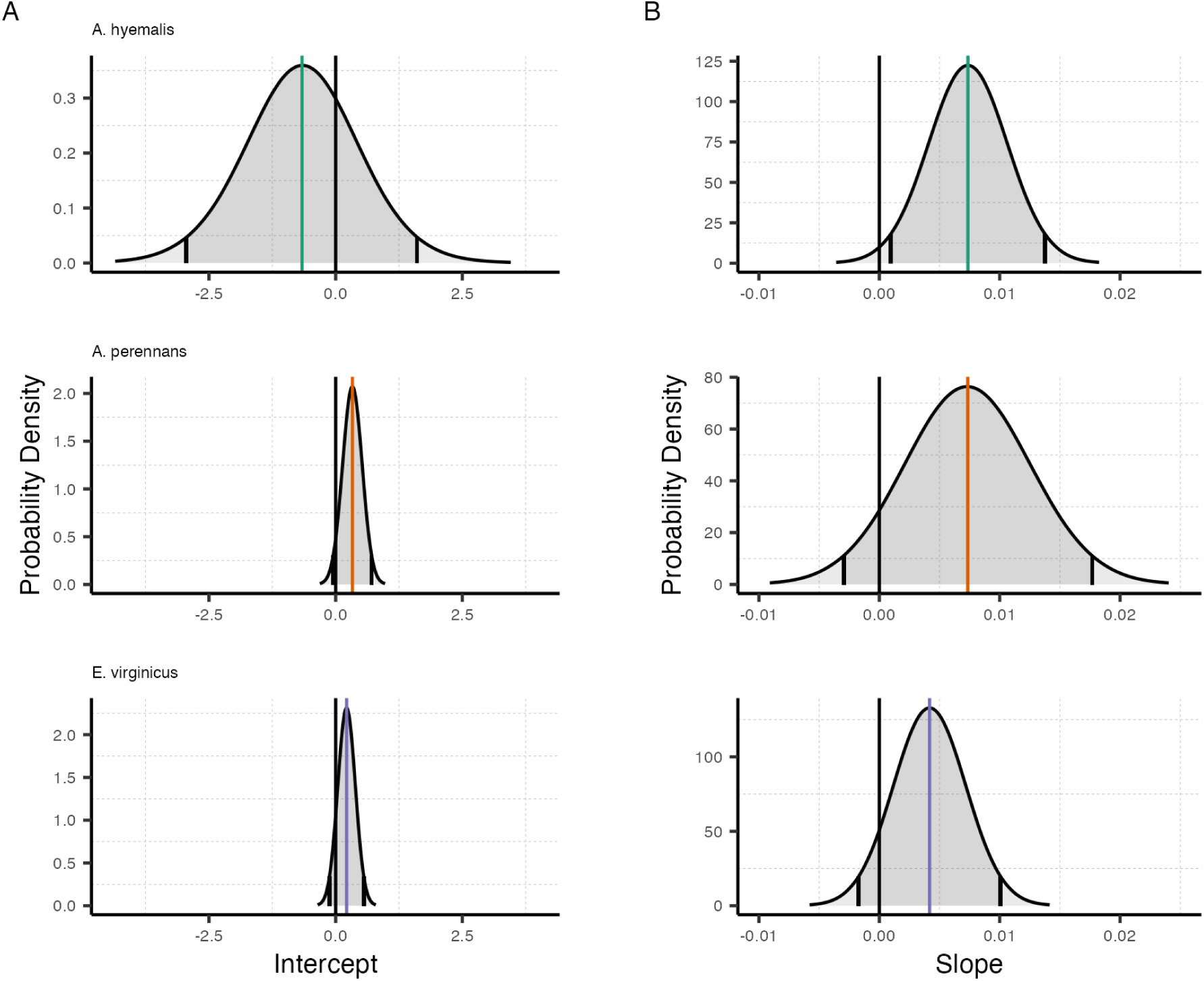
Posterior estimates of parameters describing global intercept and temporal trends from the endophyte prevalence model. Density curves show the probability density along with mean (colored line) and 95% CI (black lines) for the (A) intercept and (B) slope terms, *A* and *T* respectively from Eqn. 1. Colors represent each host species

**Figure A7:**
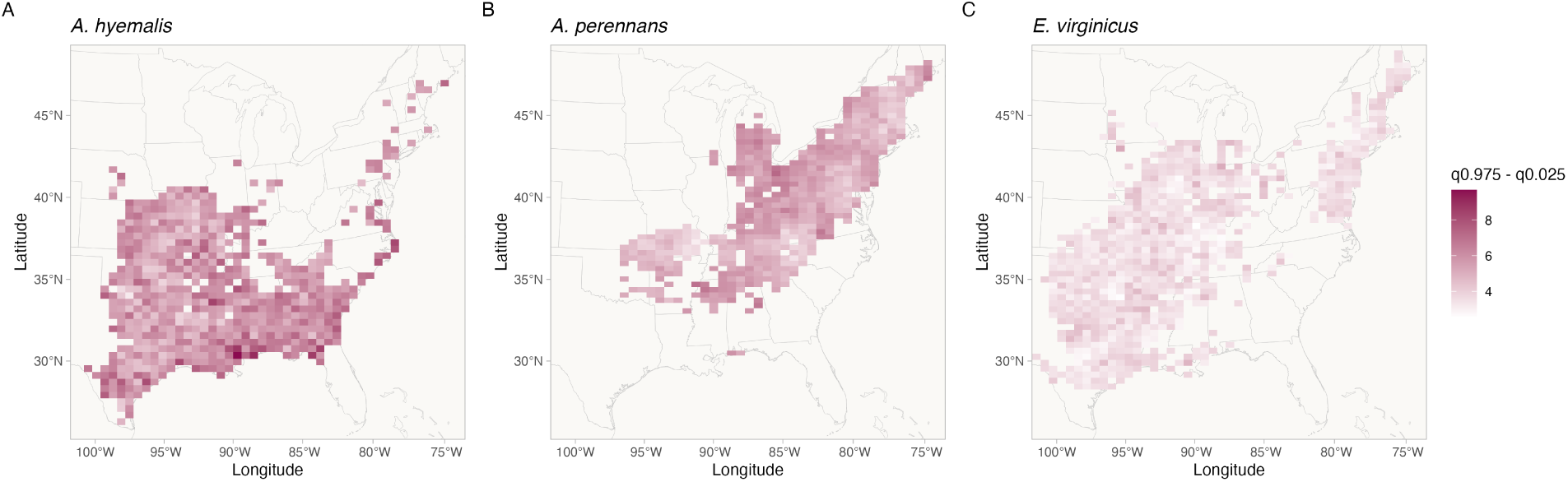
Credible interval width of temporal trends in endophyte prevalence across the distribution of each host species estimated from the endophyte prevalence model. Shading represents the range of the 95% posterior credible interval given in units of *% change in prevalence/year* for spatially varying slopes, *τ* from Eqn. 1. Map lines delineate study areas and do not necessarily depict accepted national boundaries.

**Figure A8:**
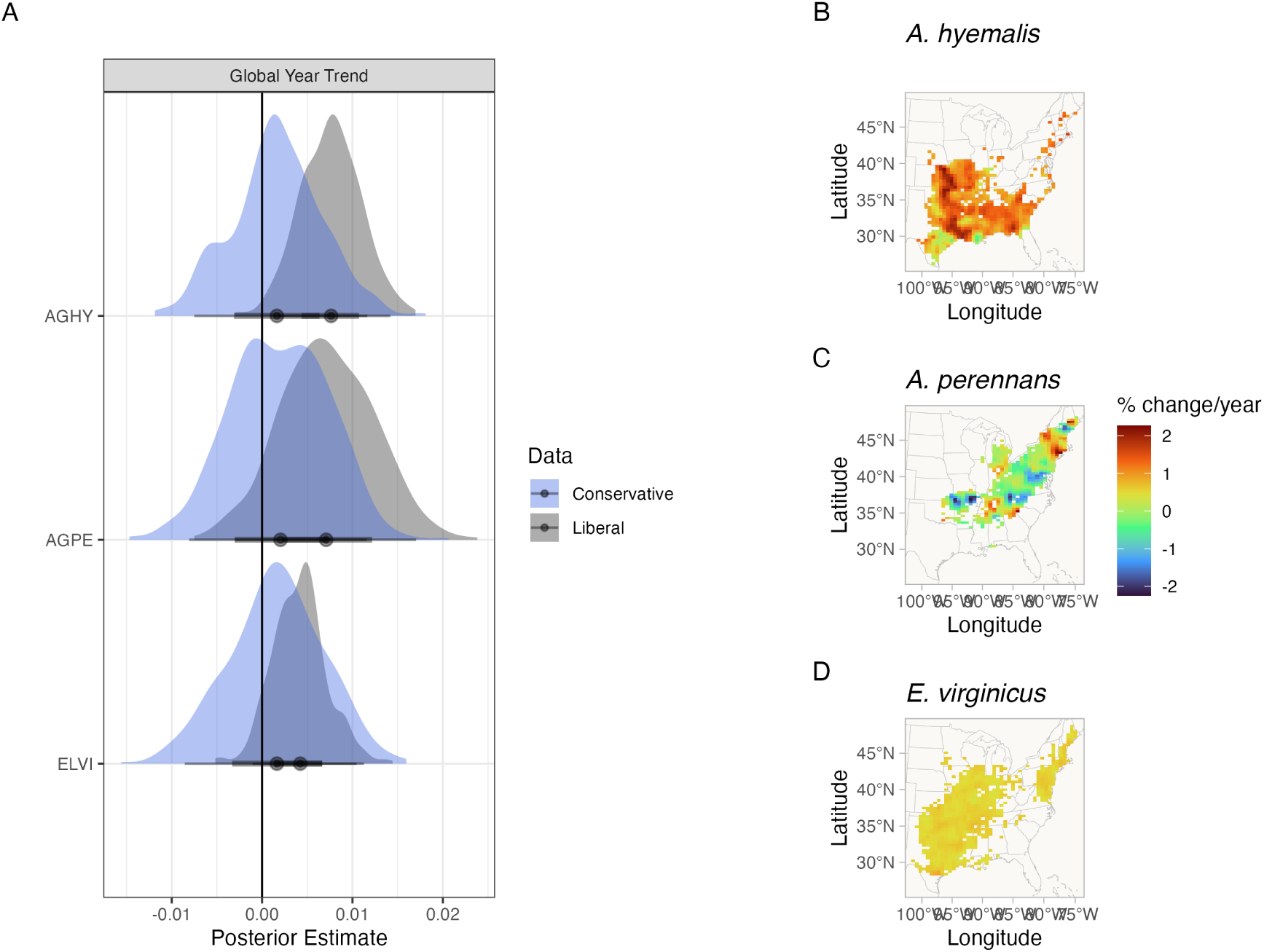
Comparison of endophyte prevalence model estimates fit to data with liberal versus conservative endophyte scores. Liberal and conservative scores document uncertainty in the endophyte identification process. Each specimen was given both a liberal and conservative scores. In cases of uncertain identification, the liberal status assumed a potential endophyte identification was more likely to be endophyte-positive while the conservative status assumed that the potential endophyte identification was less likely to be endophyte-positive. (A) Posterior estimates of global temporal trend (*T* from Eqn. 1) for the endophyte prevalence model fit to liberal scores (grey) and to conservative scores (blue). Maps show the spatially varying temporal trend estimates (*τ* from Eqn. 1) from the endophyte prevalence model fit to conservative scores for (B) *A. hyemalis*, (C) *A. perennans*, and (D) *E. virginicus*. Note that the color scale differs between this visualization and Fig. 3 that shows estimates fit using liberal endophyte scores.

**Figure A9:**
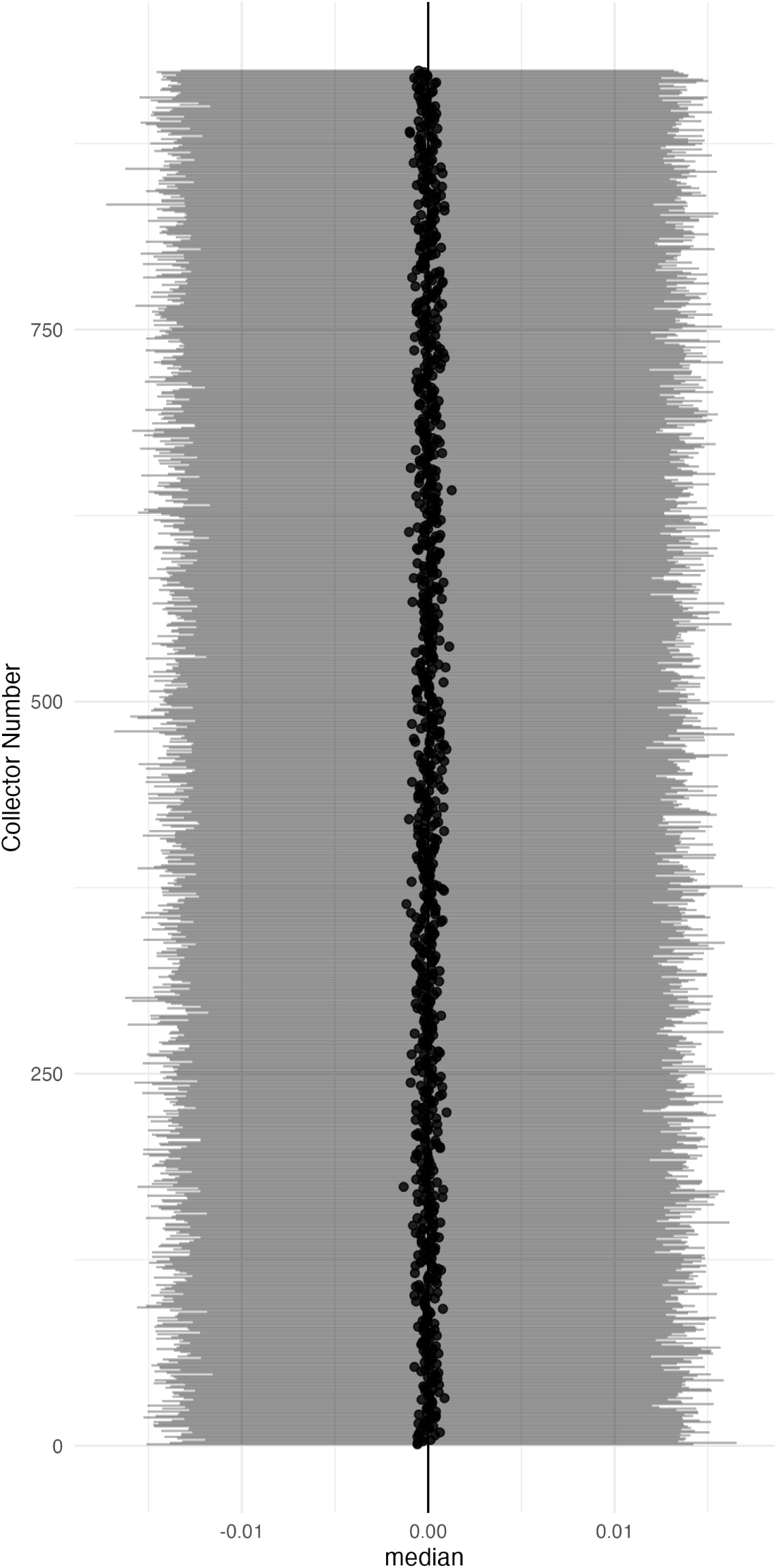
Posterior estimates of collector random effects from endophyte prevalence model. Collector random effects are denoted *χ* in Eqn. 1 and represent variance associated with researchers who collected historic herbarium specimens. Points show posterior median along with 95% CI for each of 924 individual collectors. ^59^

**Figure A10:**
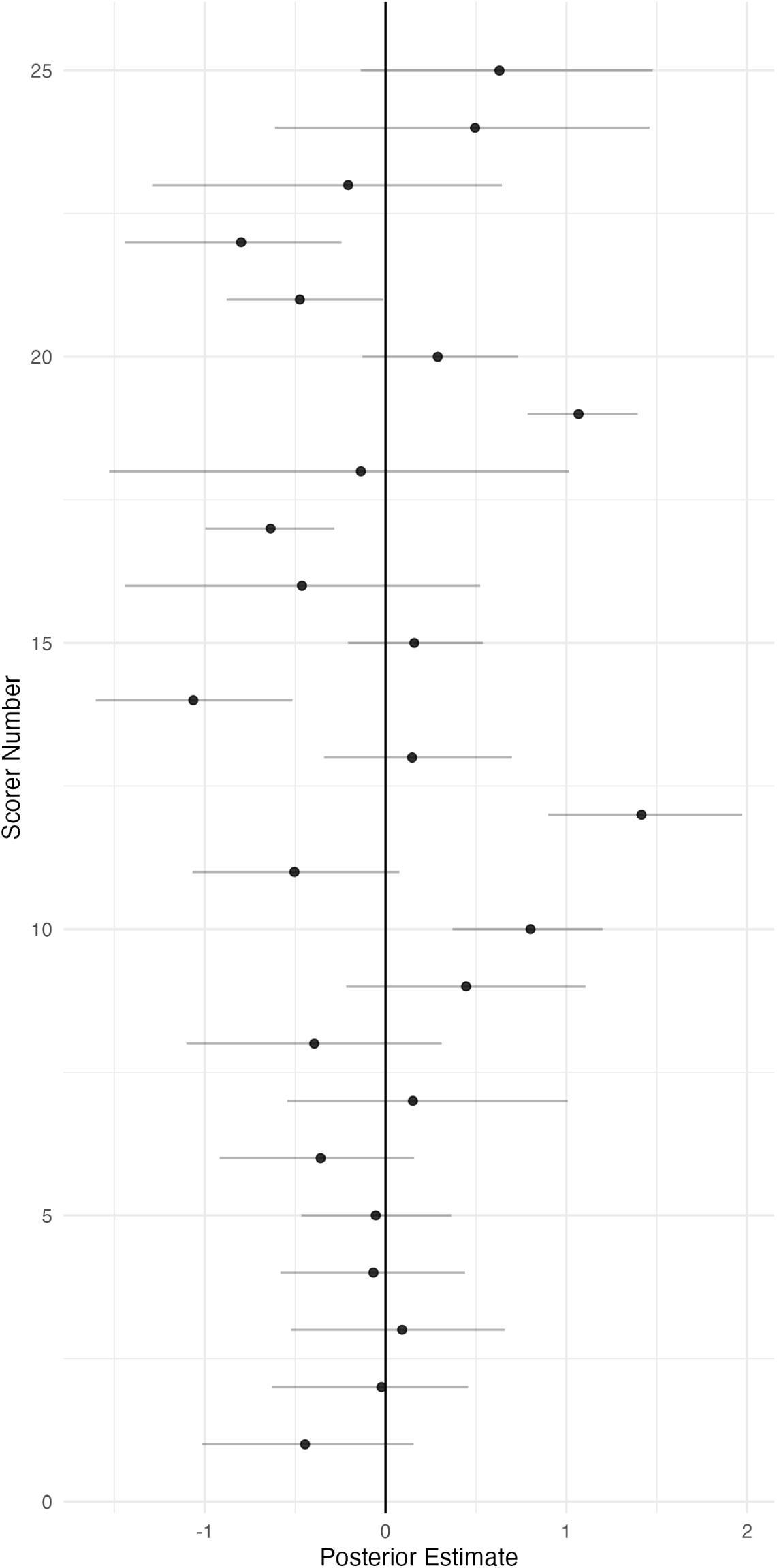
Posterior estimates of scorer random effects from endophyte prevalence model. Scorer random effects are denoted *ω* in Eqn. 1 and represent variance associated with researchers who identified *Epichloë* endophytes within herbarium specimen tissue samples. Points 60 show posterior median along with 95% CI for each of 25 individual scorers.

**Figure A11:**
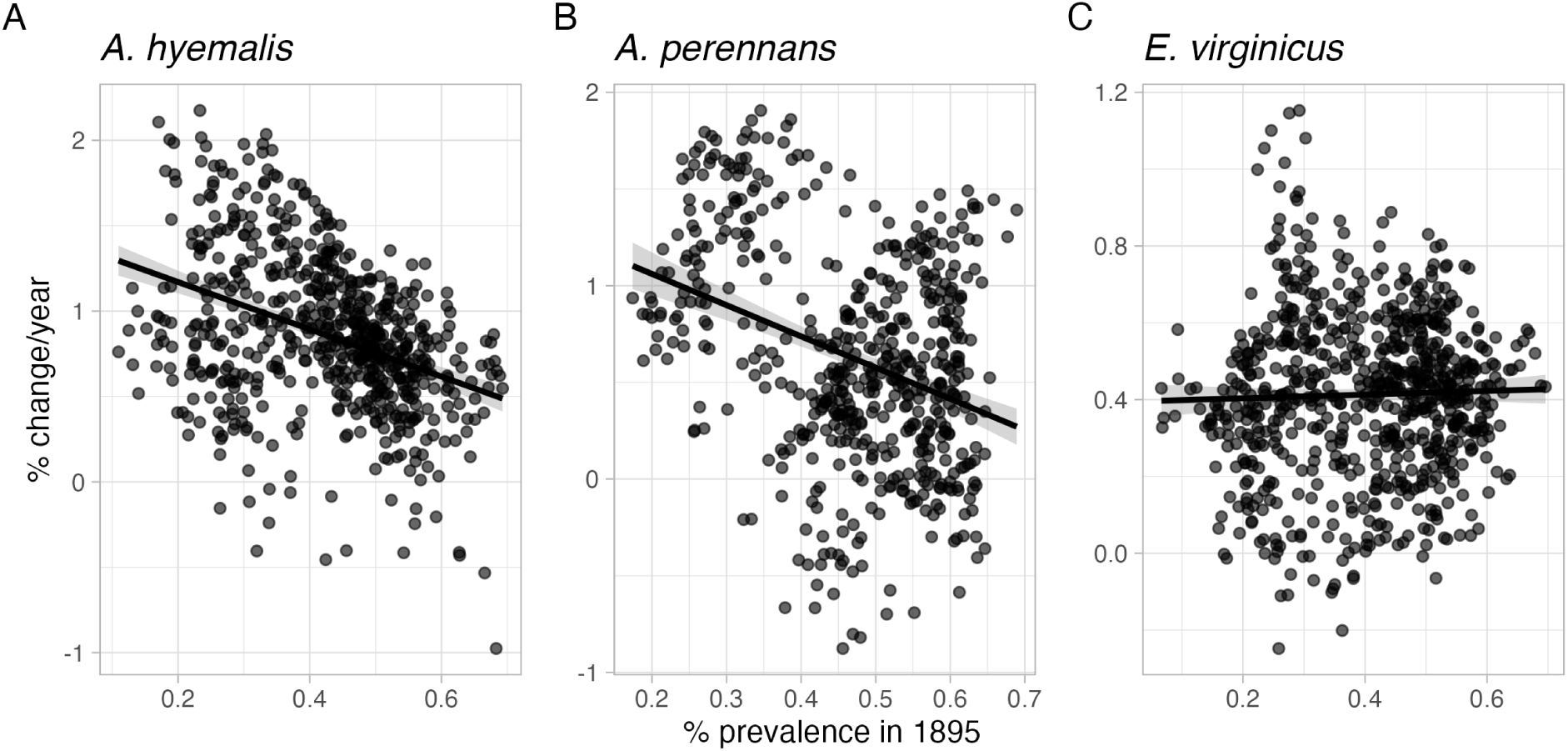
Relationship between initial prevalence and temporal trends in prevalence estimated from the endophyte prevalence model. Points show predicted posterior mean temporal trend for each species at pixels across each host distribution ((A) *A. hyemalis*, (B) *A. perennans*, and (C) *E. virginicus*). along with a linear regression and shaded ribbon showing 95% confidence interval.

**Figure A12:**
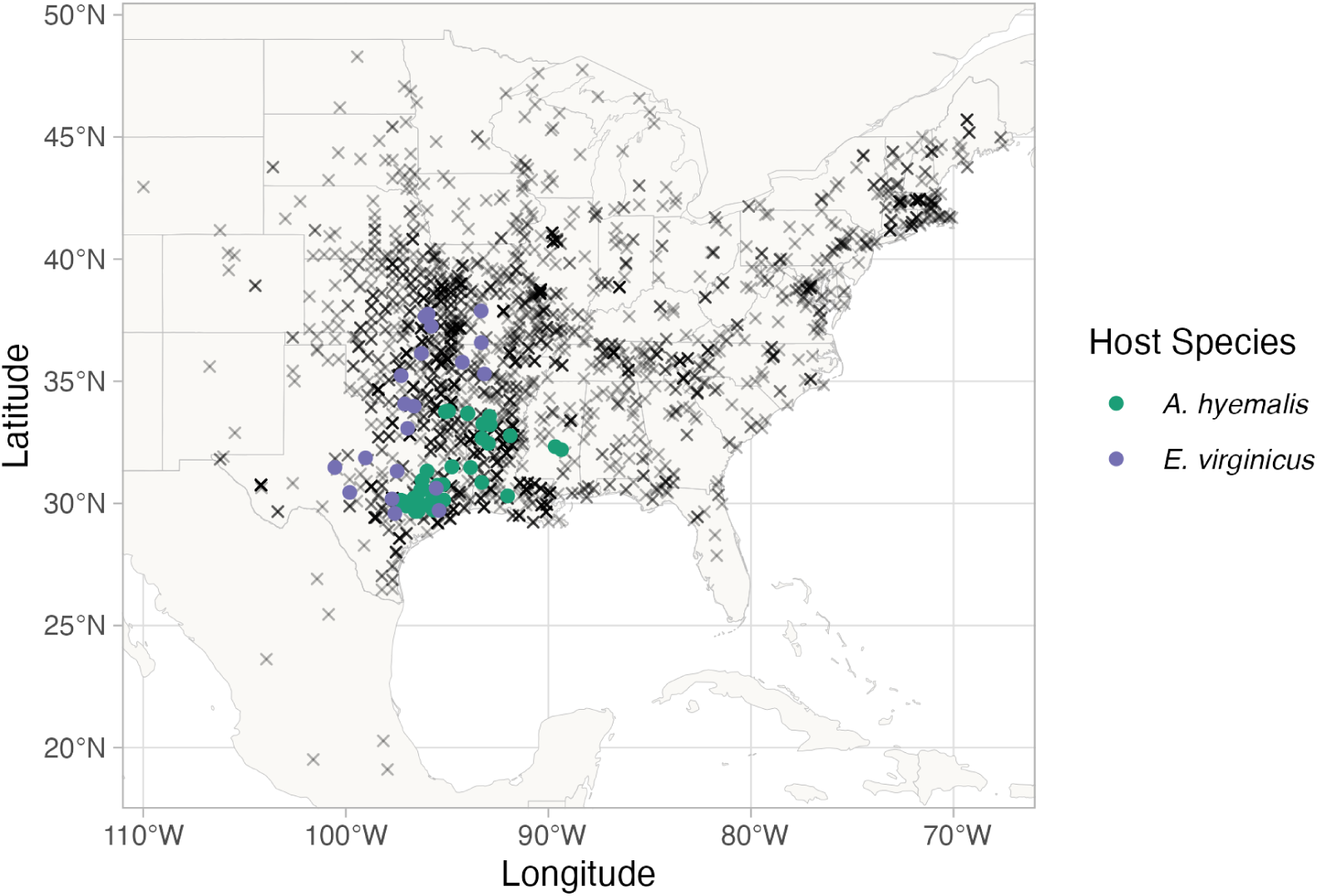
Locations of contemporary surveys of endophytes used as “test” data to evaluate predictive ability of the endophyte prevalence model. Points are locations of host populations surveyed between 2013 and 2019 for endophytes, colored by species (*A. hyemalis*: green, *E. virginicus*: purple). Black crosses show the historical herbarium collection locations used as “training” data for the endophyte prevalence model.

**Figure A13:**
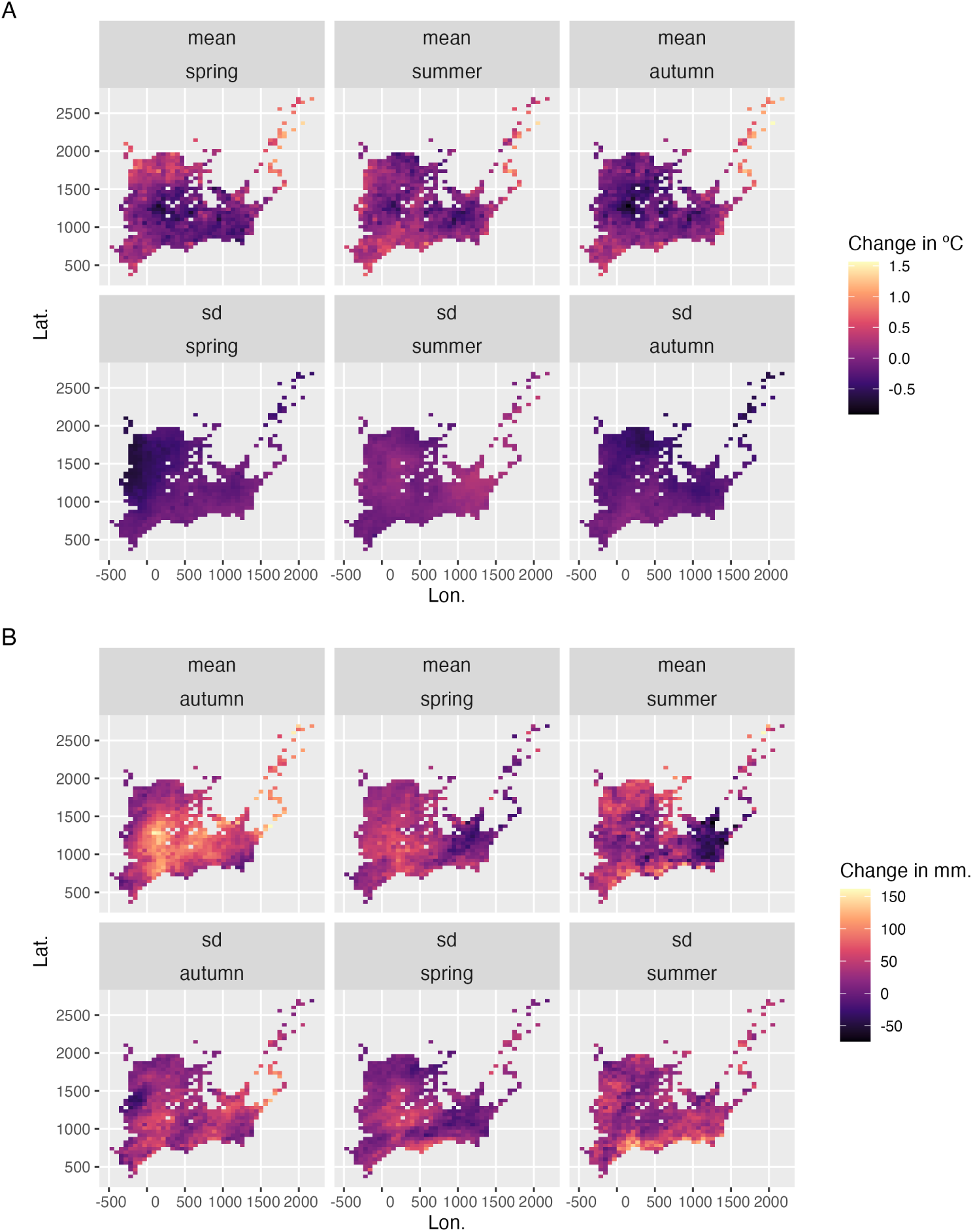
Change in seasonal climate variables between the periods 1895-1925 and 1990-2020 across the distribution of *A. hyemalis*. Color represents change in (A) seasonal temperature (*^o^*C) and (B) seasonal precipitation (mm.). Maps show pixels covering the modeled distribution of *A. hyemalis* used in *post hoc* climate regression analysis.

**Figure A14:**
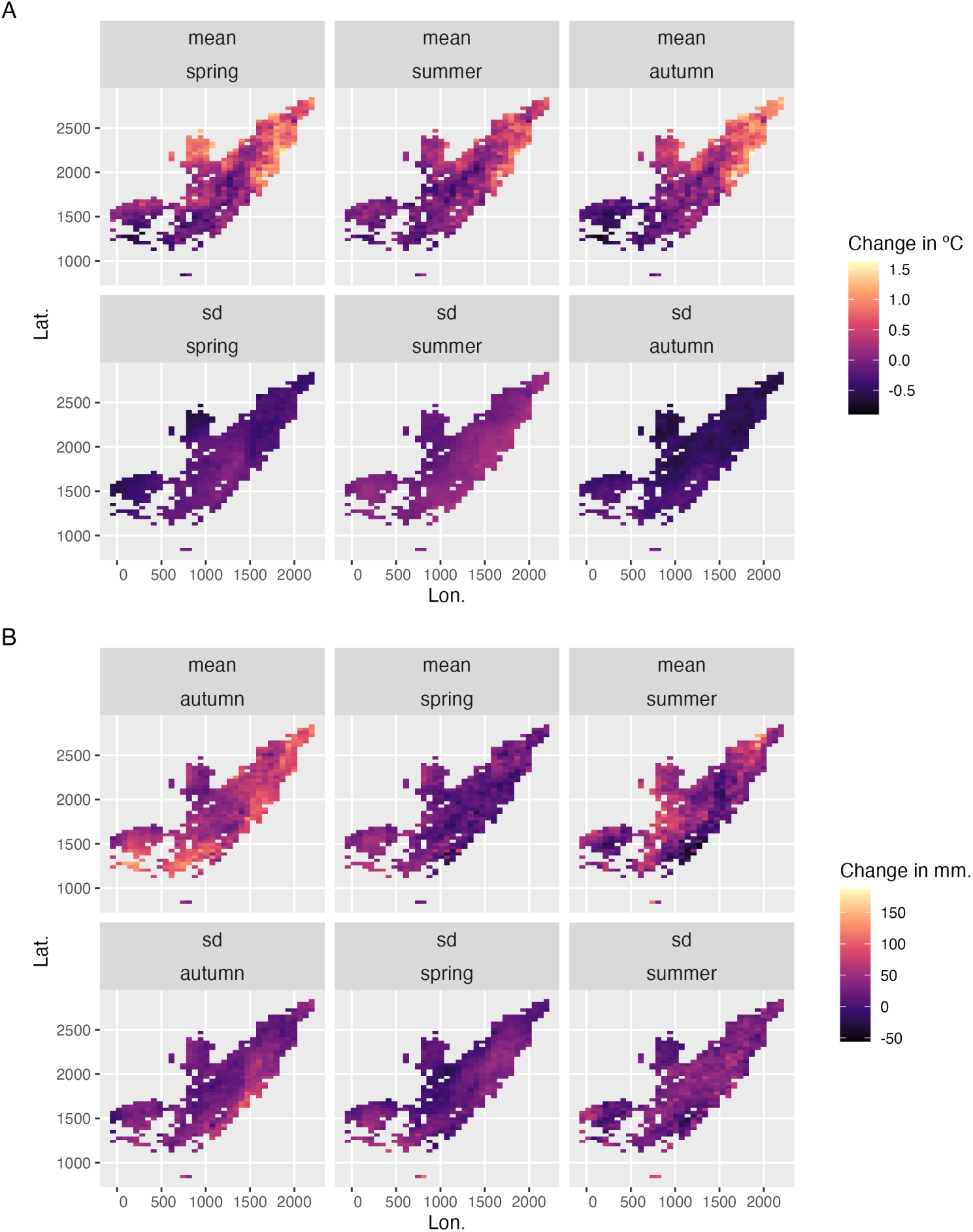
Change in seasonal climate variables between the periods 1895-1925 and 1990-2020 across the distribution of *A. perennans*. Color represents change in (A) seasonal temperature (*^o^*C) and (B) seasonal precipitation (mm.). Maps show pixels covering the modeled distribution of *A. perennans* used in *post hoc* climate regression analysis.

**Figure A15:**
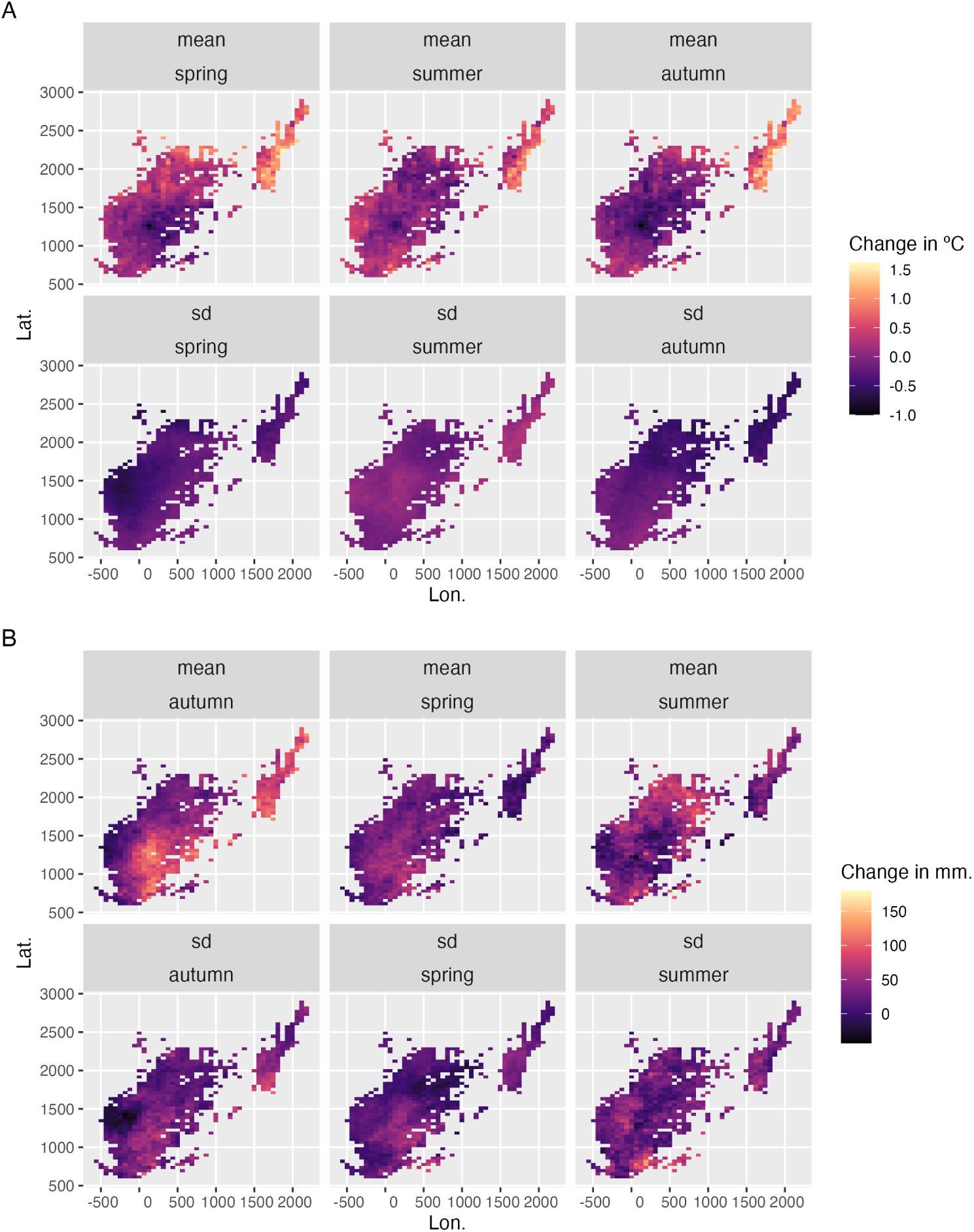
Change in seasonal climate variables between the periods 1895-1925 and 1990-2020 across the distribution of *E. virginicus*. Color represents change in (A) seasonal temperature (*^o^*C) and (B) seasonal precipitation (mm.). Maps show pixels covering the modeled distribution of *E. virginicus* used in *post hoc* climate regression analysis.

**Table A1:**
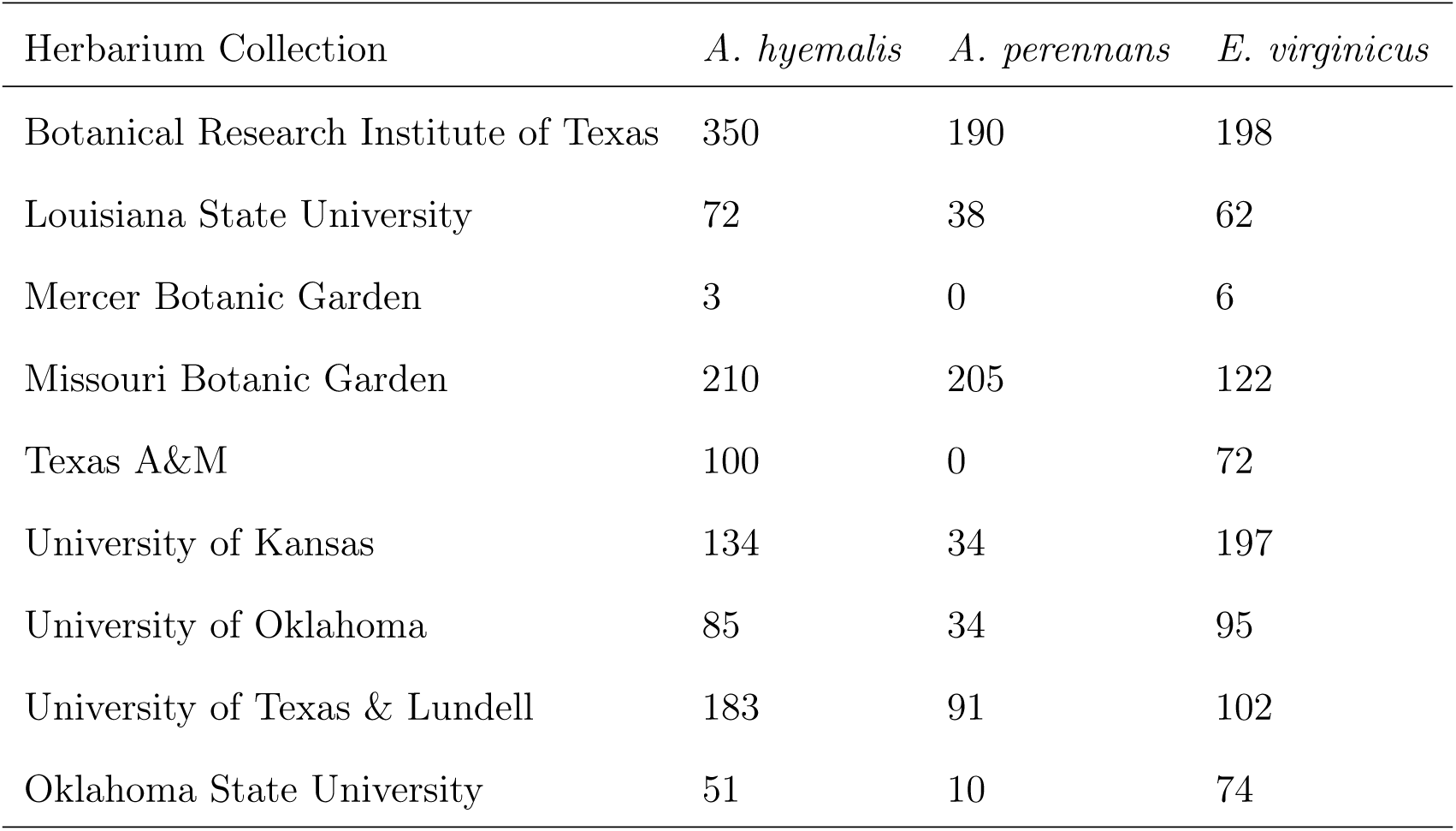
Summary of herbarium samples across collections (no. of specimens)

### Supporting Methods

#### ODMAP Protocol

##### Overview

**Model purpose**: Mapping current distribution of *Epichloë* host species.

**Target species**: *Agrostis hyemalis*, *Agrostis perennans*, and *Elymus virginicus*.

**Study area**: Eastern North America

**Spatial extent**: -125.0208, -66.47917, 24.0625, 49.9375 (xmin, xmax, ymin, ymax).

**Spatial resolution**: 0.04166667, 0.04166667 (x, y).

**Temporal extent**: 1990 to 2020.

**Boundary**: Natural.

##### Data

**Observation type**: Occurrence records from Global Biodiversity Information Facility and herbarium collection across eastern North America. We used 713 occurrences records for *Agrostis hyemalis*, 656 occurrence records for *Agrostis perennans* and 2338 for *Elymus virginicus*.

**Response data type**: occurrence record, presence-only.

**Coordinate reference system**: WGS84 coordinate reference system (EPSG:4326 code)

**Climatic data**: raster data extracted from PRISM; 30-year normal mean and standard deviation of temperature and of precipitation for three four-month seasons within the year (Spring: January, February, March, April; Summer: May; June, July, August; Autumn: September, October, November, December).

##### Model

**Model assumption**: We assumed that the target species are at equilibrium with their environment.

**Algorithms**: Maximum entropy (maxent)

**Workflow**: We described the workflow in the methods section of the manuscript.

**Software**: All statistics were performed using Maxent 3.3.4 and R4.3.1 with packages terra, usdm, spThin and dismo.

**Code availability**: Available through this link: https://github.com/joshuacfowler/EndoHerbarium

**Data availability**: Data was accessed through open-source R packages *rgbif*. *A. hyemalis* (GBIF.Org, 2025a), *A. perennans* (GBIF.Org, 2025b), *E. virginicus* (GBIF.Org, 2025c)

##### Assessment

We used AUC to test model performance.

##### Prediction

We predicted the probability of presence of the host species as a binary maps (presence or absence)

#### Mesh and Prior Sensitivity Analysis

To test the influence that the triangulation mesh and choice of priors has on results, we compared model results across a range of meshes and priors. We re-ran our model for the mesh used in main body of the text (Fig. A2), which we refer to as the “standard mesh”, and with a mesh with smaller minimum vertices (finer mesh). Finer scale meshes increase computation time. For each of these meshes, we ran the model with a range of priors defining the spatial range of our spatial random effects: 342km (the prior used for presented results), as well as ranges five times smaller (68 km) and five times larger (1714 km). We found generally that these choices did not alter the direction of model predictions, but did influence the associated uncertainty and magnitude of some effects.

For overall temporal trends, we found that models with differing priors predicted consistently positive relationships over time (Fig. A16).

**Figure A16:**
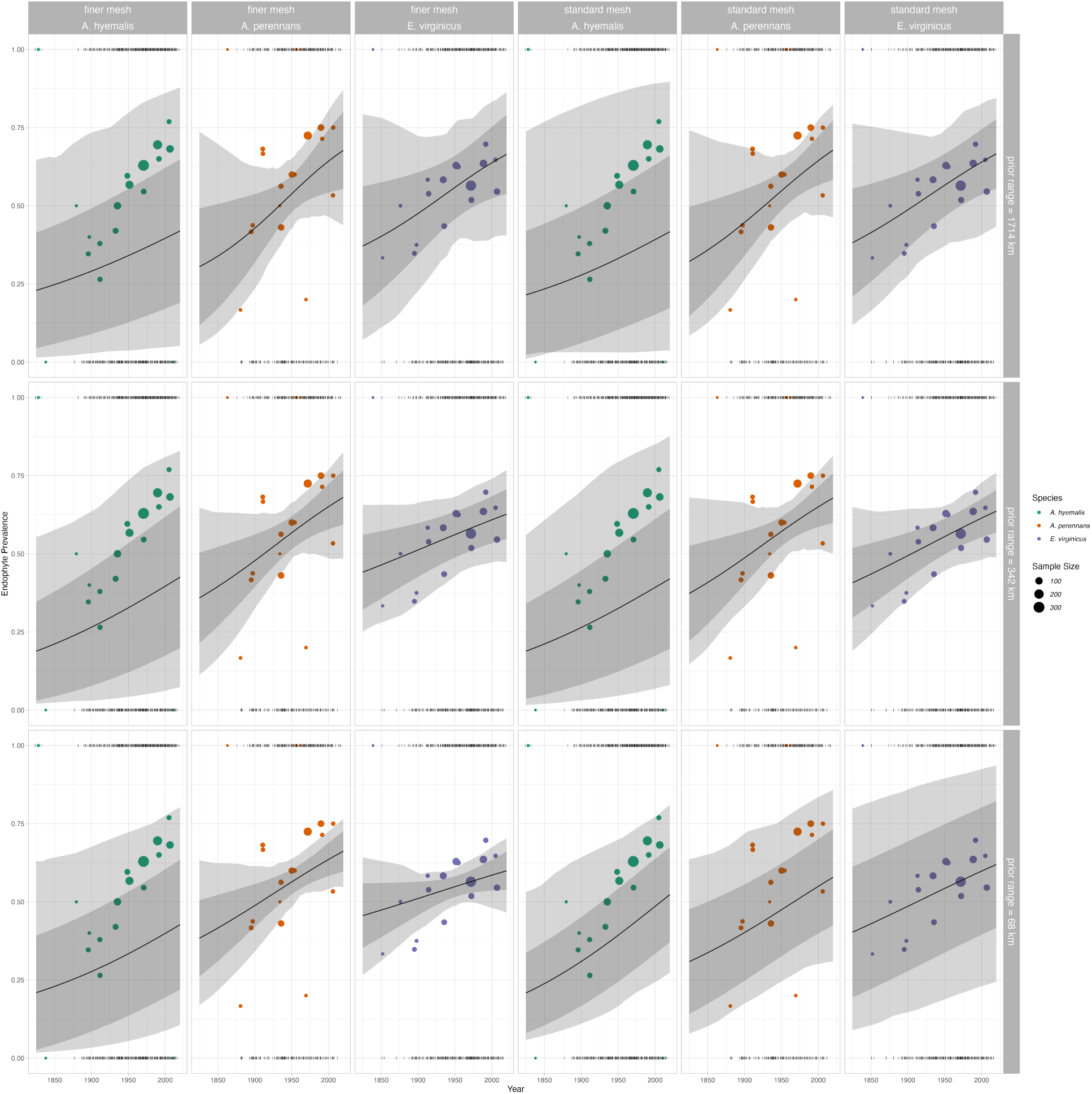
Overall trend in endophyte prevalence evaluated for endophyte prevalence models with different range priors on spatially structured random effects, and for two different triangulation meshes. Data used in model fitting is the same across all panels and as in the main text. Note that these plots, as compared to Fig. 2 in main text, show mean trends and do not incorporate variance associated with collector and scorer random effects.

For spatially-varying temporal trends, we found that models with different priors predicted consistent spatial patterns in temporal trends, although the range of this prediction varied depending on the prior and mesh (Fig. A17 - A18). One noteworthy result of this analysis is that combinations of prior choice and mesh can introduce instability in model fitting. This is evident in A17 panel B and A18 panel B, where the prior range is smaller than the minimum vertex length of the mesh. Model fitting takes an extended time period and the model struggles to identify variation across space. Results with a set of prior ranges (Fig. A17 - A and C; Fig. A18 - A and C) result in models that estimate trends across space of the same direction and order of magnitude, although the “smoothness” of these predictions vary.

**Figure A17:**
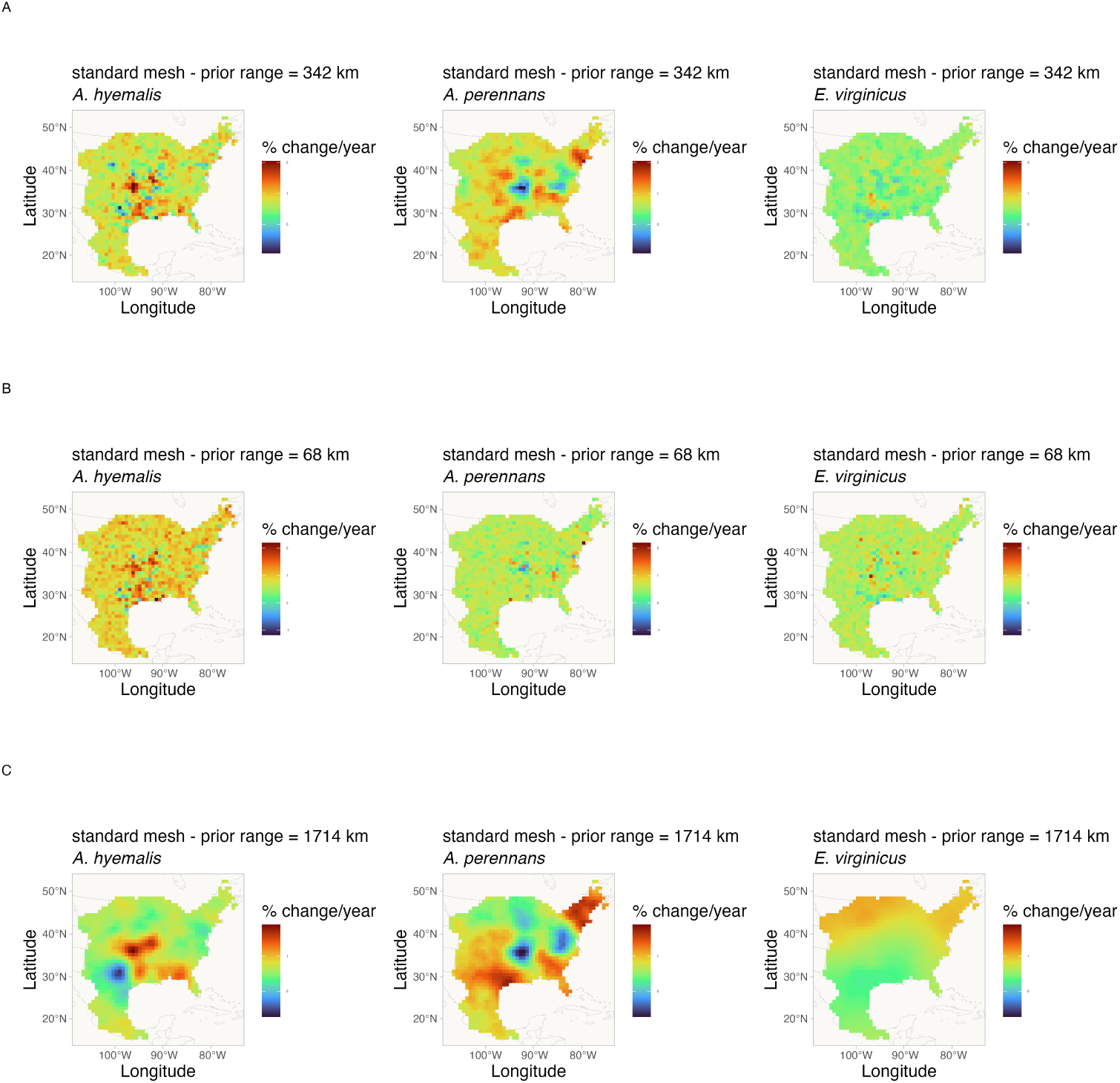
Spatially-varying trends in endophyte prevalence evaluated for the endophyte prevalence model with different range priors on spatially structured random effects, and for the “standard” mesh. Data used in model fitting is the same across all panels and as in the main text. Shading indicates the magnitude and direction of predicted trends for each of three host species for each of three prior ranges (rows A-C). Note that each plot has an individual scale bar.

**Figure A18:**
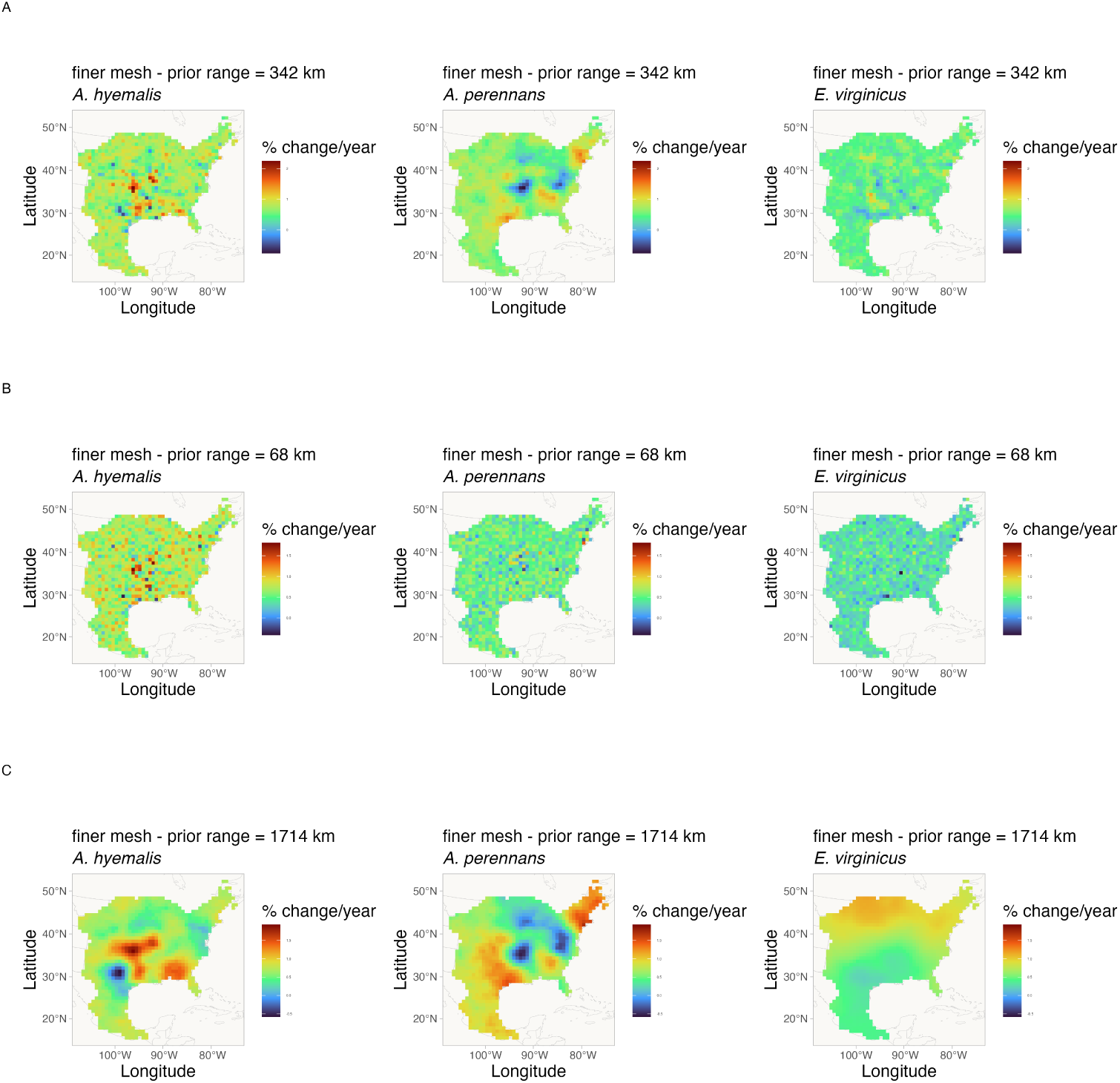
Spatially-varying trends in endophyte prevalence evaluated for the endophyte prevalence model with different range priors on spatially structured random effects, and for the “finer” mesh. Data used in model fitting is the same across all panels and as in the main text. Shading indicates the magnitude and direction of predicted trends for each of three host species for each of three prior ranges (rows A-C). Note that each plot has an individual scale bar.

#### Spatially-biased Sample Size Simulation Analysis

To examine how data that is unevenly distributed across host distributions may influence interpretation of spatially-varying coefficients, we performed a simulation analysis. Our focal species, *Agrostis hyemalis*, *Agrostis perennans*, and *Elymus virginicus*, are widely distributed grasses across the eastern United States that host *Epichloë* fungal endophytes. For logistical reasons, our sampling visits to herbaria focused on herbaria in the central southern U.S., which resulted in unevenly distributed data across each host species’ range. This is particularly noteable for *Agrostis perennans* which has the most northern distribution and relatively fewer total collected specimens compared to the other focal species. Thus, a significant portion in the northeast of this species’ range is relatively sparsely sampled. Our analysis presented in the main text identified this region as having strong increase in endophyte prevalence. Future visits to herbaria with regional focuses in the Northeastern US would certainly garner new specimens that could provide valuable insights into shifting host and symbiont distributions.

##### Simulation of spatially-biased symbiont occurence data

We simulated datasets with varying levels of missing-ness to examine how this missing-ness influenced the estimation of spatially-varying trend estimates. We first generated 300 data points for each of three hypothetical species at random positions across an area approximating the scale of our focal data. Each data point was randomly assigned a year of collection across 200 years. We then simulated data from a Bernoulli process with trends alternating across nine regions (Fig. A19) in a 3X3 grid pattern. This grid pattern was intended to create a complex spatial layout of trends, where trends were either an increase of 1% per year or a decrease of 1% per year.

**Figure A19:**
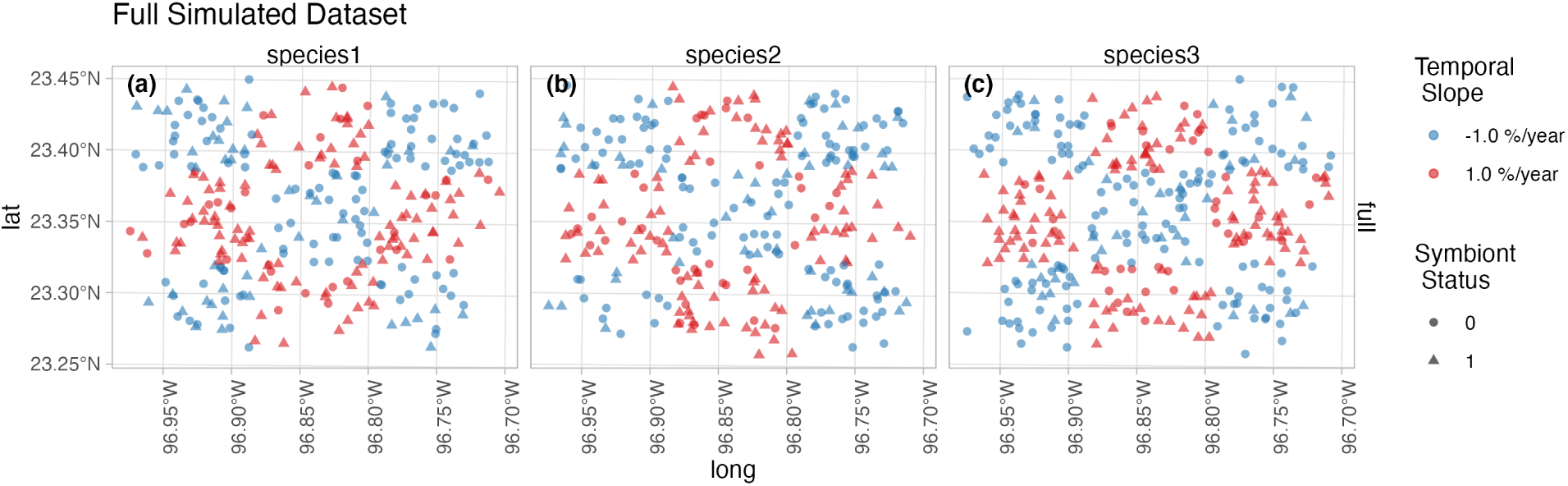
Full simulated dataset of symbiotic association with spatially-varying temporal trends. Color indicates the slope parameter used to simulate trends in endophyte status across nine “regions” for three species. Data are assigned collection years across a period of 200 years. Shape indicates the presence (1) or absence (0) of a symbiont.

From this full data, we generated six additional datasets with missing-ness in the northeast region of the simulated data for hypothetical species 2. The data remained the same for Species 1 and for species 3 across all datasets. For these six datasets, we removed data points at random in six ways: 0% of datapoints in northeast region, 0% of recent datapoints, only 20% of datapoints, only 20% of recent datapoints, only 50% of datapoints, and only 50% of recent datapoints (Fig. A20). We define the datapoints as part of the recent time period if they occur later than the median year. The result is 6 scenarios exploring degrees of spatial and temporal bias.

**Figure A20:**
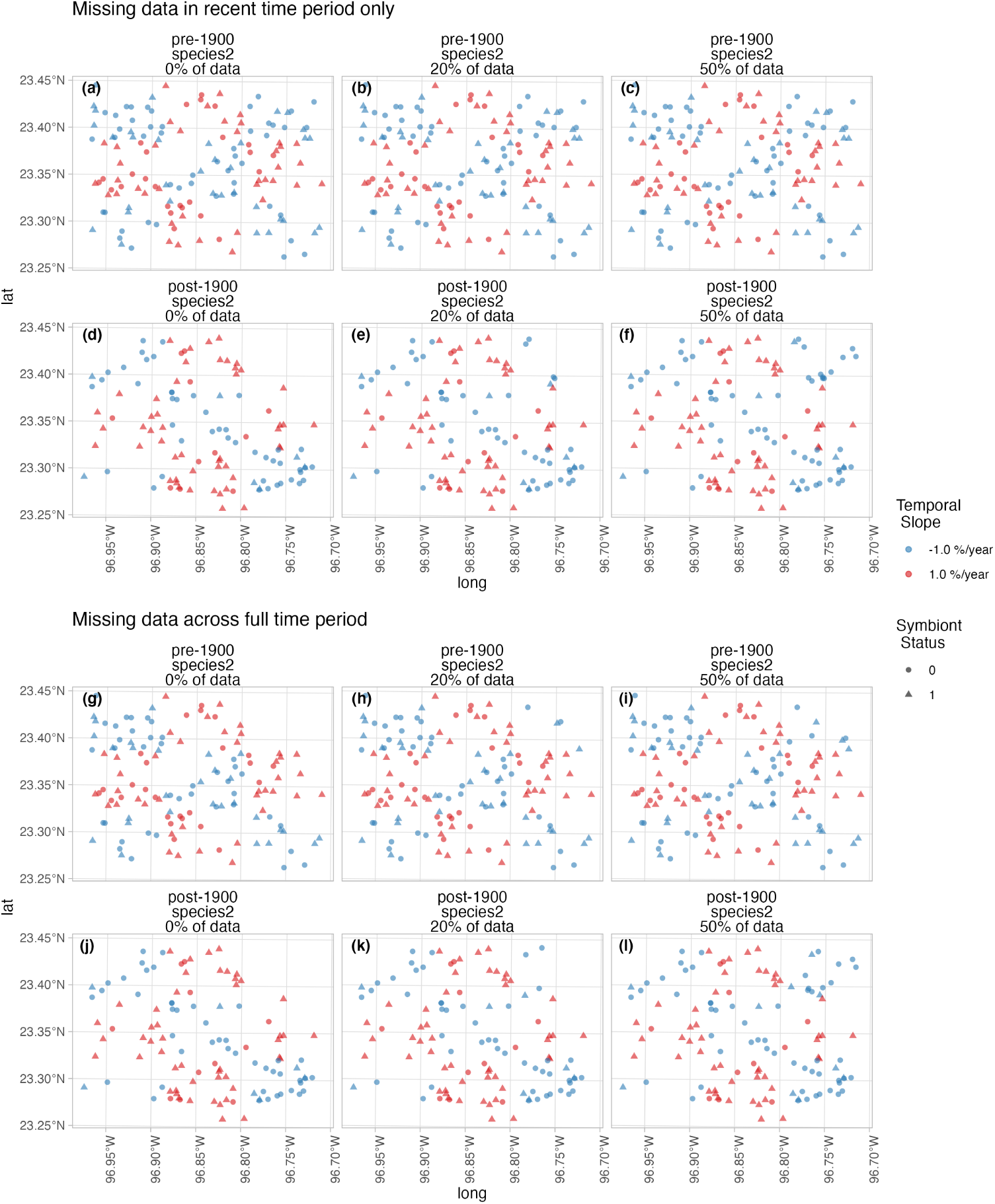
Six simulated datasets representing scenarios of spatially-baised missingness for Species 2. Missingness was imposed in the northeast region for six scenarios: 0% of recent datapoints available (a,d); only 20% of recent datapoints (b,e); only 50% of recent datapoints (c,f); 0% of datapoints across the full time period available (g,j); only 20% of datapoints across the full time period (h,k); and only 50% of datapoints across the full time period(i,l). Missingness was imposed only for hypothetical Species 2; Species 1 and 3 remain as in Figure A19. Color indicates the slope parameter used to simulate trends in endophyte status across 9 regions in a 3×3 grid. Shape indicates the presence (1) or absence (0) of a symbiont.

##### Statistical analysis

We analyzed each dataset with a model given by Eqn. A1 similar in construction to that used in our central analysis.

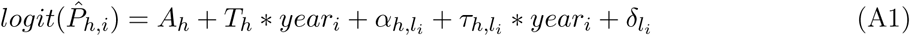

Where symbiont presence/absence of the *i^th^* specimen (*P_h,i_*) was modeled as a Bernoulli response variable with expected probability of symbiont occurrence *P̂_h,i_* for each host species *h*. We modeled *P̂_h,i_* as a linear function of intercept *A_h_* and slope *T_h_* defining the global trend in endophyte prevalence specific to each host species as well as with spatially-varying intercepts *α_h,l__i_* and slopes *τ_h,l__i_* associated with location (*l_i_*, the unique latitude-longitude combination of the *i*th observation). Similar to the SVC model of our central analysis (Eqn. 1), we estimated a shared variance term with the spatially-dependent random effect *δ_l__i_* , intended to account for residual spatial variation. However in this analysis we omit i.i.d.-random effects terms associated with collector and scorer identity (*χ_c__i_* and *ω_s__i_* in Eqn. 1) for the sake of simplicity.

##### Influence of spatially-biased sampling on model interpretation

Our analysis of the full simulated data shows that our model is suitably flexible to capture complex spatial patterns in temporal trends (Fig. A21 a-c). Beyond this, the model also qualitatively captures the spatial patterns in temporal trends even with large amounts of data missingness (i.e missing up to 80% of the datapoints (Fig. A21 p-r)).

While this analysis is not an exhaustive examination of the influence of sampling bias on our results for several reasons (including not examining how different strengths in temporal trends, different spatial arrangments of missing-ness influence model estimates, or different sample sizes influence results), it demonstrates that the spatially-varying modelling framework implemented in INLA we employ can suitably recover regional trends even with significant spatially-bias within data collection, and further the analysis is likely robust to temporally-structured bias (missing data within recent collection period). Future work could more fully explore the scenarios that cause this ability to break down. We expect this simulation reflects what may be a common scenario for research investigating global change using natural history specimens. Collection effort by trained taxonomists and professional collectors peaked in the past, and collections contain relatively fewer modern specimens in many regions. Additionally, most global change research necessarily involves accessing many specimens across collections. Research efforts such as ours will be unable to access every specimen from all possible collections. Ongoing digitization efforts will make it possible to more clearly assess how much data is missing from a particular study compared to the actual holdings of natural history collections, but ultimately, the decision of what data and collections to include is a question of sample size and study design.

**Figure A21:**
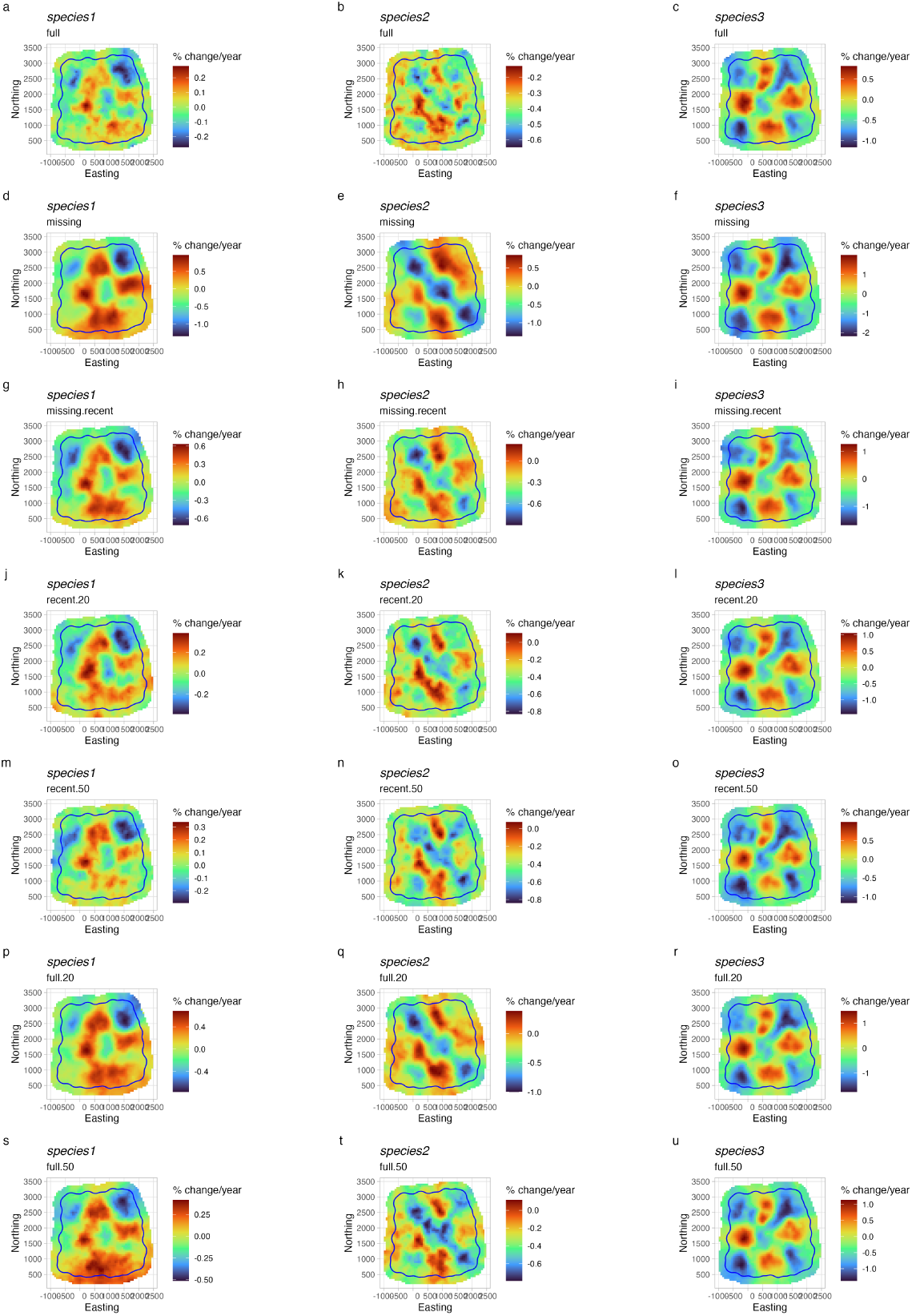
Mean predicted spatially-varying trend in symbiont prevalence across datasets with different levels of missingness. Color indicates the estimated mean temporal trend within each pixel across the simulated data. Panels show estimates for models fit to different levels of missing data for species 2 in the northeast region ((a-c) the full dataset, (d-f) missing all datapoints across entire temporal period, (g-i) missing all datapoints only during the recent period, (j-l) missing 80% of the datapoints only during the recent period, (m-o) missing 50% of the datapoints only during the recent period, (p-r) missing 80% of the datapoints across the entire temporal period, (s-u) missing 50% of the datapoints across the entire temporal period). The mesh boundary that bounds the “full” simulated dataset is plotted in each panel.

